# A genome-scale CRISPR screen reveals PRMT1 as a critical regulator of androgen receptor signaling in prostate cancer

**DOI:** 10.1101/2020.06.17.156034

**Authors:** Stephen Tang, Nebiyou Y. Metaferia, Marina F. Nogueira, Maya K. Gelbard, Sarah Abou Alaiwi, Ji-Heui Seo, Justin H. Hwang, Craig A. Strathdee, Sylvan C. Baca, Jiao Li, Shatha AbuHammad, Xiaoyang Zhang, John G. Doench, William C. Hahn, David Y. Takeda, Matthew L. Freedman, Peter S. Choi, Srinivas R. Viswanathan

## Abstract

Androgen receptor (AR) signaling is the central driver of prostate cancer across disease states. While androgen deprivation therapy (ADT) is effective in the initial treatment of prostate cancer, resistance to ADT or to next-generation androgen pathway inhibitors invariably arises, most commonly through re-activation of the AR axis. Thus, orthogonal approaches to inhibit AR signaling in advanced prostate cancer are essential. Here, via genome-scale CRISPR/Cas9 screening, we identify protein arginine methyltransferase 1 (PRMT1) as a critical mediator of *AR* expression and signaling. PRMT1 regulates recruitment of AR to genomic target sites and inhibition of PRMT1 impairs AR binding at lineage-specific enhancers, leading to decreased expression of key oncogenes, including *AR* itself. Additionally, AR-driven prostate cancer cells are uniquely susceptible to combined AR and PRMT1 inhibition. Our findings implicate PRMT1 as a key regulator of AR output and provide a preclinical framework for co-targeting of AR and PRMT1 in advanced prostate cancer.

## INTRODUCTION

There is currently no curative therapy for castration-resistant prostate cancer (CRPC), the lethal phase of prostate cancer that arises when the disease progresses on androgen deprivation therapy (ADT). Although CRPC may be driven by either androgen-dependent or androgen-independent resistance mechanisms, the former, which involve re-activation of androgen receptor (AR) transcriptional activity, are far more common (Watson et al., 2015). Mechanisms leading to sustained AR signaling in CRPC include *AR* gene (Visakorpi et al., 1995) and/or *AR* enhancer (Dessel et al., 2019; Quigley et al., 2018; Viswanathan et al., 2018a) amplification, activating point mutations in *AR* (Taplin et al., 1995), intra-tumoral steroid production, and synthesis of constitutively active truncated *AR* splice variants (*AR-V*s) such as *AR-V7* (Dehm et al., 2008).

Collectively, 85-90% of CRPC tumors display genomic alterations at the *AR* locus (Li et al., 2020) and a comparable percentage display heightened AR activity upon transcriptional profiling (Chen et al., 2004), indicating that robust maintenance of AR signaling output is pervasive in CRPC despite suppression of androgen levels by ADT. These observations have motivated the development of next-generation androgen pathway inhibitors such as enzalutamide and abiraterone. Although such agents have shown benefit in CRPC, resistance invariably emerges (Watson et al., 2015).

Multiple mechanisms of resistance to next-generation androgen pathway inhibitors have been described. Commonly, somatic alterations at the *AR* locus or production of *AR-V*s lead to re-activation of AR signaling (Viswanathan et al., 2018a; Takeda et al., 2018; Antonarakis et al., 2014). For example, production of *AR-V7* results from inclusion of a cryptic exon (cryptic exon 3, *CE3*) during splicing of *AR* pre-mRNA, leading to expression of a protein product that contains the N-terminal transactivation domain of AR but lacks its C-terminal ligand binding domain. In contrast to full-length AR, this truncated receptor possesses ligand-independent transactivation and repressive activities (Cato et al., 2019; Dehm et al., 2008). Other resistance mechanisms include overexpression of AR coactivators (Hwang et al., 2019; Lee et al., 2019), activation of AR bypass pathways such as glucocorticoid receptor signaling (Arora et al., 2013), or activation of AR-independent oncogenic pathways such as Myc (Han et al., 2017) or Wnt (Grasso et al., 2012; Robinson et al., 2015).

Given the compelling genetic evidence that AR signaling plays a critical role in both initial castration resistance and advanced CRPC, orthogonal approaches to modulate AR output are of significant therapeutic interest. While candidate-based approaches have previously identified a limited number of factors regulating *AR* expression (Wang et al., 2016), genome-scale functional genetic technologies can enable unbiased discovery of such modulators and can anticipate resistance mechanisms that may arise in patients. For example, a genome-scale open reading frame (ORF) screen recently implicated the transcription factor CREB5 as a mediator of enzalutamide resistance via its ability to enhance AR activity at a subset of enhancers and promoters (Hwang et al., 2019), and a genome-scale CRISPR/Cas9 screen implicated HNRNPL in regulating alternative splicing of *AR* (Fei et al., 2017).

Here, we applied genome-scale CRISPR/Cas9 genetic screening to identify key regulators of *AR*/*AR-V7* expression. We identified protein arginine methyltransferase 1 (PRMT1) as a critical mediator of *AR* expression and signaling that is required for the binding of AR to its genomic target sites. Genetic or pharmacologic targeting of PRMT1 inhibits *AR* and AR target gene expression and impairs viability in multiple cellular models of activated AR signaling. Our results provide a preclinical rationale for co-targeting of AR and PRMT1 in AR-driven CRPC.

## RESULTS

### Genome-scale CRISPR/Cas9 screening identifies regulators of *AR*/*AR-V7* expression

We sought to systematically identify regulators of *AR*/*AR-V7* expression by leveraging genome-scale genetic screening in a cellular model of CRPC. The prostate cancer cell line 22Rv1 expresses high levels of the truncated *AR* splice variant *AR-V7*, which promotes resistance to the next-generation antiandrogen enzalutamide and is required for androgen-independent growth (Cato et al., 2019; Li et al., 2013). To identify transcriptional or post-transcriptional regulators of *AR/AR-V7* expression, we first used CRISPR/Cas9 targeting with homology-directed repair to introduce a GFP-containing cassette in-frame with *CE3* in 22Rv1 cells (Figure 1A). We isolated two independent knock-in clones (22Rv1/AR-V7-GFP/Clone 6 and 22Rv1/AR-V7-GFP/Clone 9) and verified successful integration of the GFP-containing cassette in both clones (Figure S1A). As expected, GFP expression in the knock-in cell line was suppressed upon silencing of either total *AR* or *AR-V7* mRNA, but not upon selective silencing of full-length *AR* (*AR-FL*) mRNA (Figures S1B and S1C).

**Figure 1.**
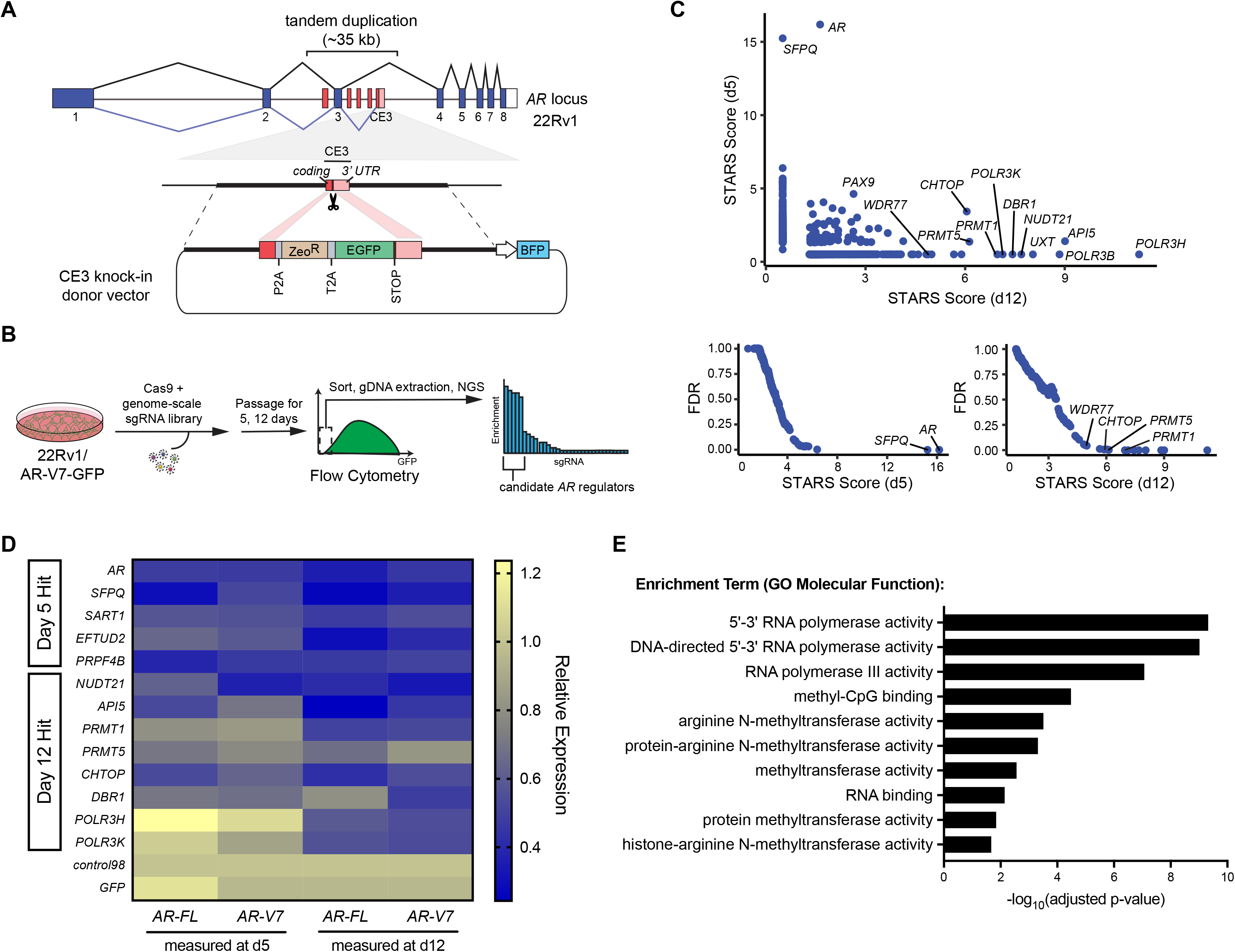
Genome-scale CRISPR/Cas9 screen identifies regulators of *AR*/*AR-V7* expression. (A) Schematic of 22Rv1/AR-V7-GFP reporter cell line. CRISPR/Cas9 editing and homology-directed repair were used to insert a GFP-containing cassette directly upstream of the cryptic exon 3 (CE3) stop codon in 22Rv1 cells. (B) Schematic of the CRISPR/Cas9 screening strategy used to identify regulators of *AR*/*AR-V7* expression in 22Rv1/AR-V7-GFP cells. (C) Screen hits, plotted by STARS score on day 5 or day 12 after library transduction, determined by enrichment of sgRNAs in the sorted GFP-negative population as compared with the starting library pool. *Top*: Scatterplot of STARS scores for screen hits on day 5 versus day 12. For plotting purposes, hits that scored at only one timepoint were assigned a STARS score of 0.5 for the timepoint that they were not enriched. *Bottom*: Plots of false discovery rate (FDR) versus STARS score of hits from day 5 (left) or day 12 (right) timepoints. Selected high-scoring hits are labeled. (D) Arrayed validation of screen hits by RT-qPCR in parental 22Rv1 cells. Heatmap shows relative *AR-FL* and *AR-V7* expression in 22Rv1 cells at the indicated timepoints after knockout of selected screen hits. mRNA levels are normalized to a control sgRNA (control98). Data are presented as the mean of n = 4 replicates. (E) Enrichment analysis showing gene ontology (GO) terms significantly enriched among screen hits scoring on either day 5 or day 12 with FDR < 0.25.

Having established a faithful cellular reporter of *AR/AR-V7* expression, we next proceeded to genome-scale CRISPR/Cas9 screening. We transduced 22Rv1/AR-V7-GFP/Clone 6 with a Cas9 expression vector followed by a pooled library of 76,441 barcoded sgRNAs targeting 19,114 unique genes. At either 5 or 12 days after library transduction, GFP-negative cells were sorted by flow cytometry and gDNA was extracted for detection of sgRNA barcodes by next-generation sequencing (Figure 1B). Cells were assayed at two timepoints, given that genes in lineage-essential pathways might selectively score at an earlier time point and drop out at a later time point, while genes with long protein half-lives or delayed phenotypic effects might score only at a later time point. Candidate regulators of *AR/AR-V7* expression were identified based on enrichment of sgRNAs compared with the starting library pool. Using a false discovery rate (FDR) threshold of FDR < 0.25, we identified 27 significant hits at the day 5 timepoint and 19 significant hits at the day 12 timepoint (Figure 1C and Tables S1 and S2). We validated 13 selected top hits from our screen in an arrayed fashion in both Clone 6 and Clone 9 cells (Figure S1D), and observed excellent concordance between the independent clones across all hits tested (Figure S1E).

Given the design of our reporter cell line, hits from the screen could represent selective post-transcriptional regulators of *AR-V7* expression, or transcriptional/post-transcriptional regulators of *AR-V7* as well as other *AR* species. To distinguish between these possibilities, we knocked out 13 hits from our primary screen in parental 22Rv1 cells and measured the effects on *AR-FL* and *AR-V7* transcript expression (Figure 1D). Interestingly, knockout of certain genes (e.g. *NUDT21*, *DBR1*) showed more pronounced effects on *AR-V7* expression than *AR-FL* expression, nominating these genes as selective regulators of *AR-V7*. In contrast, knockout of most tested genes resulted in comparable decreases in both *AR-FL* and *AR-V7* expression, implicating these candidates in the regulation of total, rather than isoform-specific, *AR* expression. We also observed that certain hits scored preferentially at either the early (e.g. *AR*, *SFPQ*) or late (*PRMT1, POLR3H, POLR3K*) timepoints (Figure 1D).

To determine whether hits from our screen converged on common biological processes, we performed gene ontology enrichment analysis on all 46 genes that scored (FDR < 0.25) at either the day 5 or day 12 timepoint. In addition to several terms associated with RNA processing and mRNA splicing, we observed a striking enrichment for gene ontology terms related to arginine methyltransferase activity (Figure 1E). This was driven by highly scoring hits encoding protein arginine methyltransferases (PRMT1, PRMT5), proteins found in complex with PRMTs (CHTOP, WDR77) (Takai et al., 2014), and PRMT substrates such as SFPQ (Snijders et al., 2015) and AR itself (Mounir et al., 2016).

PRMTs comprise a family of nine enzymes that catalyze the deposition of methyl groups on arginine residues of substrate proteins, resulting in diverse biological consequences. These enzymes are divided into two predominant subclasses based on the methylation pattern that they deposit; type I PRMTs (e.g. PRMT1, PRMT4, PRMT6) place asymmetric dimethyl (ADMA) marks, while type II PRMTs (e.g. PRMT5, PRMT7) place symmetric dimethyl (SDMA) marks. Of the multiple hits from our screen involved with protein arginine methylation, we elected to further characterize *PRMT1* for the following reasons. First, PRMT1-mediated arginine dimethylation of histone 4 results in a transcriptionally activating mark (H4R3me2a) that may be associated with prostate cancer recurrence (Seligson et al., 2005; Strahl et al., 2001; Wang, 2001). Second, PRMT1 has been reported to associate with nuclear hormone receptors, including AR, and to function as a transcriptional coactivator (Koh et al., 2001; Wang, 2001). And third, PRMT1 is amenable to inhibition by small molecules, including multiple tool compounds and a type I PRMT inhibitor currently in clinical development (Eram et al., 2016; Fedoriw et al., 2019; Fong et al., 2019; Yan et al., 2014).

### PRMT1 regulates *AR* expression and signaling in advanced prostate cancer

We next sought to extend our observation that PRMT1 regulates *AR/AR-V7* expression in several prostate cancer cell lines. The 22Rv1 and VCaP cell lines co-express *AR-FL* and *AR-V7* while the LNCaP cell line exclusively expresses *AR-FL*. *PRMT1* knockdown reduced *AR-FL* and *AR-V7* expression in 22Rv1 and VCaP cells and *AR*-*FL* expression in LNCaP cells, as determined by RT-qPCR (Figure 2A). Similar results were observed at the protein level, as well as in both 22Rv1/AR-V7-GFP knock-in clones (Figures S2A-S2C).

**Figure 2.**
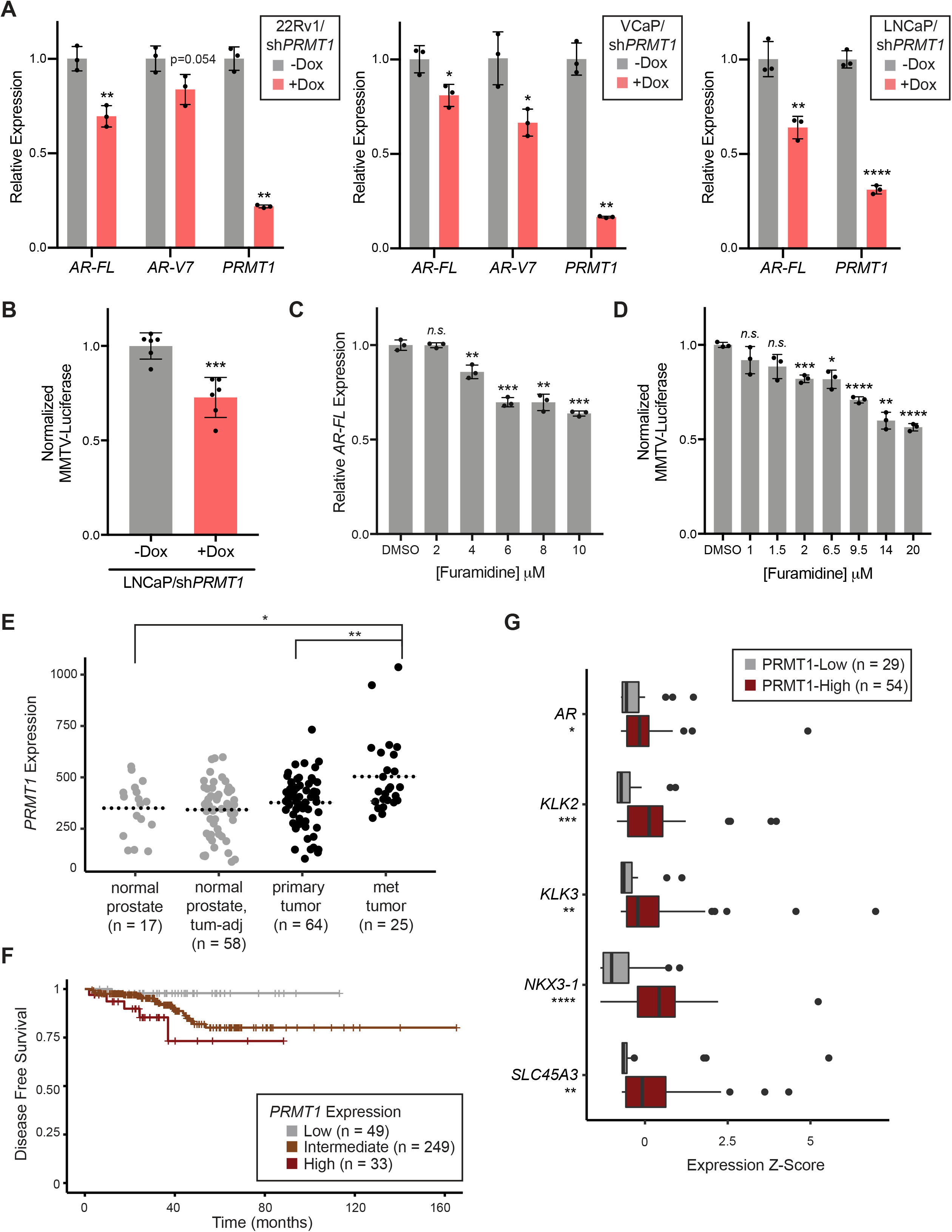
PRMT1 regulates *AR* expression and AR signaling in advanced prostate cancer. (A) Relative *AR-FL*, *AR-V7*, and *PRMT1* expression, as assessed by RT-qPCR, with or without *PRMT1* knockdown by dox-inducible shRNA in the prostate cancer cell lines 22Rv1, VCaP, and LNCaP. Expression is shown relative to no dox. Error bars represent mean ± SD, n = 3 replicates. (B) Relative luciferase activity upon *PRMT1* knockdown in LNCaP cells transduced with an androgen-responsive MMTV-Luciferase reporter. Luciferase activity is normalized to cell viability for each condition and shown relative to no dox. Error bars represent mean ± SD, n = 6 replicates. (C) Relative *AR-FL* expression in LNCaP cells after treatment with the PRMT1 inhibitor furamidine at the indicated doses. Expression is shown relative to DMSO. Error bars represent mean ± SD, n = 3 replicates. (D) Relative MMTV-Luciferase activity in LNCaP cells upon treatment with furamidine at the indicated doses. Luciferase activity is normalized to cell viability at each concentration and shown relative to DMSO. Error bars represent mean ± SD, n = 3 replicates. (E) *PRMT1* expression in normal prostate tissue, tumor-adjacent normal tissue, primary prostate tumors, and metastatic tumors from a published dataset (Chandran et al., 2007). The *y*-axis represents signal intensity values from an oligonucleotide microarray probe against *PRMT1*. Statistical significance was determined by Mann-Whitney U test. (F) Kaplan-Meier plot showing disease-free survival after prostatectomy among prostate cancer patients with low, intermediate, or high *PRMT1* expression in a published dataset (Hoadley et al., 2018). The survival distributions of the three groups are significantly different (log-rank test, p = 0.03). (G) Relative expression of *AR* and target genes *KLK2*, *KLK3*, *SLC45A3*, and *NKX3-1* in published mRNA expression data (Abida et al., 2019) from castration-resistant prostate cancer tumors with low or high *PRMT1* expression. Statistical significance was determined by Mann-Whitney U test. For A-D, statistical significance was determined by Student’s t-test. *p < 0.05, **p < 0.01, ***p < 0.001, ****p < 0.0001.

Given that *PRMT1* suppression has a comparable effect on both *AR-V7* and *AR-FL* expression, we hypothesized that PRMT1 may regulate *AR* mRNA transcription. We performed a pulse-chase assay using an ethynyl uridine (EU) label to examine the kinetics of *AR* mRNA synthesis and decay. We found that the rate of *AR* transcription decreased upon *PRMT1* knockdown while the rate of *AR* mRNA decay was not significantly affected, suggesting that PRMT1 regulates *AR* expression by modulating the production, rather than the stability, of *AR* mRNA (Figure S2D). Next, to determine the consequences of *PRMT1* suppression on AR signaling, we employed a reporter construct expressing luciferase under the control of an androgen-responsive mouse mammary tumor virus (MMTV) promoter. *PRMT1* knockdown in LNCaP cells transduced with the MMTV-Luciferase reporter led to a significant decrease in luciferase activity (Figure 2B). Finally, we assessed whether the decreases in *AR* expression and signaling observed with genetic targeting of *PRMT1* could be recapitulated with pharmacologic PRMT1 inhibition. Indeed, upon treatment of LNCaP cells with the PRMT1 inhibitor furamidine, we observed dose-dependent decreases in both *AR* expression and MMTV-Luciferase activity (Figures 2C and 2D).

Next, we interrogated published prostate cancer datasets to examine the relationship between PRMT1 and disease aggressiveness in human tumor samples. We observed a stepwise increase in *PRMT1* expression with prostate cancer disease stage, with higher expression in metastatic tumors than in primary tumors and normal prostate tissue (Figure 2E) (Chandran et al., 2007). Higher *PRMT1* expression was also associated with shorter recurrence-free survival after prostatectomy (Figure 2F) (Hoadley et al., 2018). Given that AR signaling is commonly sustained in CRPC tumors despite castrate testosterone levels, and that PRMT1 has been shown to enhance nuclear receptor target gene transactivation (Koh et al., 2001), we next asked whether *PRMT1* expression correlates with AR signaling output in CRPC. Strikingly, we observed elevated expression of *AR* and its canonical target genes *KLK2*, *KLK3*, *NKX3-1*, and *SLC45A3* in CRPC tumors with high *PRMT1* expression compared to tumors with low *PRMT1* expression (Figure 2G) (Abida et al., 2019). Altogether, these data suggest that PRMT1 may play an important role in the progression of prostate cancer to a hormone-refractory state through activation of *AR* expression and signaling.

### PRMT1 modulates the expression and splicing of AR target genes

PRMT1 has been reported to elicit diverse effects on gene regulation via modification of substrate proteins, including histones and RNA binding proteins (Li et al., 2010; Zhang et al., 2015). To evaluate the transcriptomic consequences of *PRMT1* suppression, we performed RNA-sequencing (RNA-seq) on LNCaP cells in the presence or absence of *PRMT1* knockdown. We observed global changes in gene expression upon *PRMT1* knockdown, with more genes downregulated than upregulated (1,721 versus 568 genes at |log_2_FC| > 0.5 and q < 0.05) (Table S3). Strikingly, several canonical AR target genes, including *KLK2*, *KLK3*, *NKX3-1*, and *SLC45A3*, were among the most significantly downregulated genes with *PRMT1* knockdown. A previous study of genome-wide AR binding sites in normal and tumor prostate tissue identified a set of 324 genes that are upregulated in tumor relative to normal prostate tissue and are proximal to tumor-specific AR binding sites (t-ARBSs) (Pomerantz et al., 2015). We observed significant enrichment of this gene set among genes downregulated upon *PRMT1* knockdown (Figure 3A), suggesting that PRMT1 is a critical regulator of the tumor-specific AR cistrome. Consistent with this notion, enrichment analysis revealed that the genes most downregulated upon *PRMT1* knockdown were enriched for AR targets as well as genes normally co-expressed with several transcription factors critical to prostate tumorigenesis, including NKX3-1, HOXB13, FOXA1, and ERG (Figure 3B).

**Figure 3.**
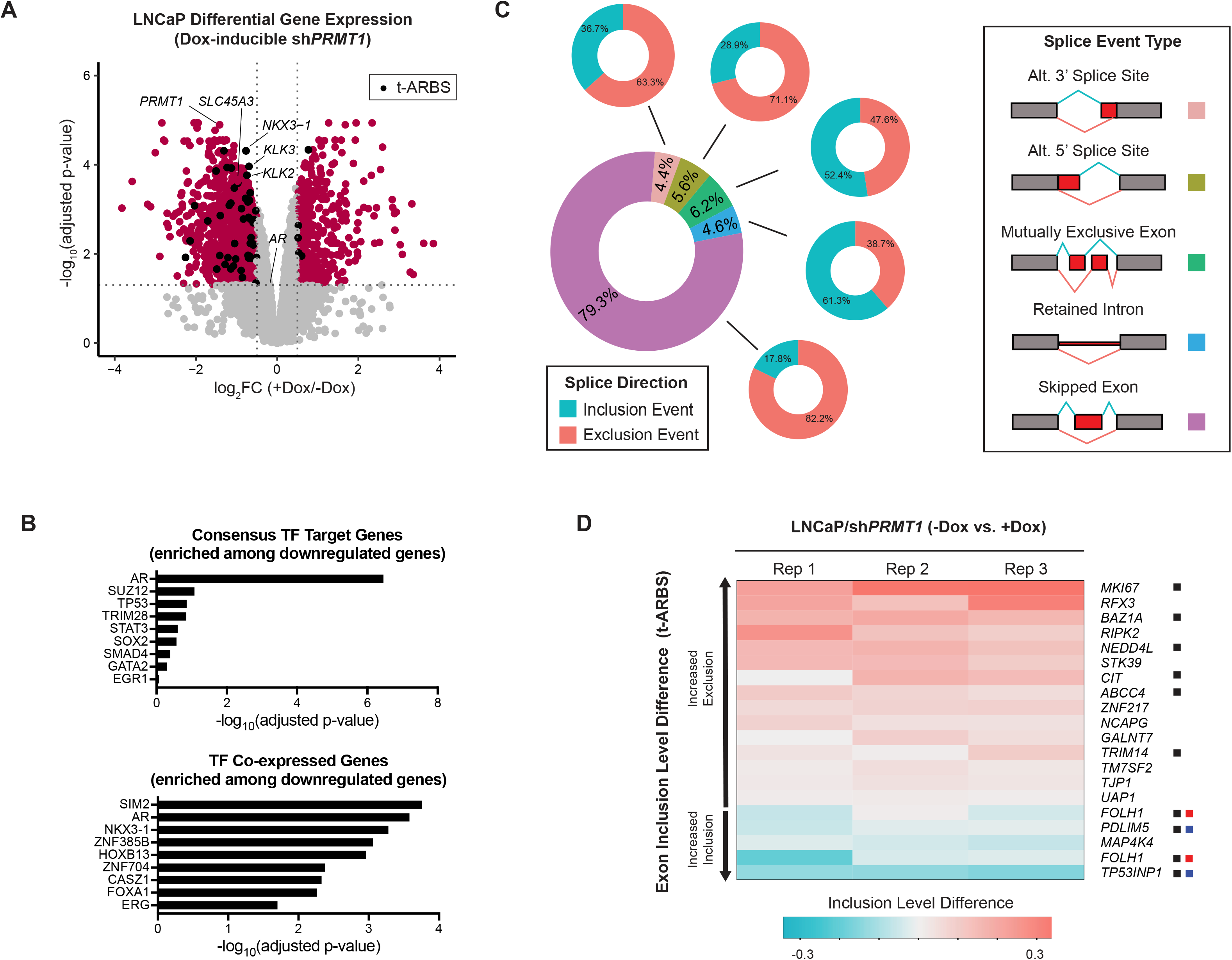
*PRMT1* suppression globally perturbs expression and splicing of AR target genes. (A) Volcano plot showing differentially expressed genes in LNCaP cells upon *PRMT1* knockdown by dox-inducible shRNA, as assessed by transcriptome sequencing. Genes meeting significance and differential expression thresholds of adjusted p-value < 0.05 and |log_2_FC| > 0.5 are colored in red. Of these, genes proximal to tumor-specific AR binding sites (t-ARBSs) (Pomerantz et al., 2015), are colored in black and are significantly enriched among downregulated genes (Fisher’s exact test, p = 0.002). *AR*, *PRMT1*, and four selected canonical AR target genes are labeled. n = 3 replicates were used in each condition. (B) Enrichment analysis of genes downregulated upon knockdown of *PRMT1*. The top 100 downregulated genes are enriched for targets of the transcription factors listed in the top panel and are highly co-expressed with transcription factors listed in the bottom panel. (C) Global alterations in splicing patterns observed in the setting of *PRMT1* knockdown. *Left*: Central donut plot shows the proportion of each splice event type among all differential splicing events observed; peripheral donut plots show the proportion of inclusion and exclusion events within each splice event type. *Right*: Schematic of different splice event classes. ‘Inclusion’ events are shown in blue and ‘exclusion’ events are shown in red. (D) Heatmap of inclusion level difference (ILD) among differentially utilized exons in t-ARBS genes upon *PRMT1* knockdown. Exons are rank-ordered by ILD. Positive ILD values (red) correspond to increased relative exon skipping while negative ILD values (blue) correspond to increased relative exon inclusion upon *PRMT1* knockdown. The corresponding gene to which each exon belongs is indicated at right. Significant gene-level expression differences are indicated with a black box (adjusted p-value < 0.05), red box (log_2_FC < −0.5), and/or blue box (log_2_FC > 0.5).

Given the reported role of PRMTs in modifying RNA binding proteins, which in turn can influence pre-mRNA splicing, we next sought to understand how *PRMT1* suppression affects splicing patterns in prostate cancer cells. Differential alternative splicing analysis revealed that *PRMT1* knockdown predominantly affected exon usage compared to other splice event classes (79% differential exon usage), with a majority of differential exon usage events representing exon exclusion (82% exclusion) (Figures 3C and S3A; Table S4). Interestingly, we noted that of the 728 unique genes exhibiting differential exon usage, significant gene-level expression differences (q < 0.05) were observed in only 25% (185/728). Thus, the majority of splicing changes upon *PRMT1* knockdown occurred independently of changes in level of gene expression, suggesting effects on factors that directly regulate pre-mRNA splicing independent of gene transcription.

Among the top excluded (skipped) exons, we observed an enrichment of AR target genes (Figure S3B), suggesting that in addition to downregulation of genes in proximity to t-ARBSs, *PRMT1* knockdown might also induce splicing alterations that affect AR signaling. Overall, we identified 20 differential exon inclusion events affecting t-ARBS genes, the majority of which represented excluded exons (75% excluded; Figure 3D). For example, the most pronounced exon exclusion event was found in the gene *MKI67*, which encodes a cell proliferation marker, Ki-67, commonly used in prognostic evaluation of prostate and other cancers. Ki-67 exists in cells in two predominant splice isoforms distinguished by the inclusion or exclusion of exon 7, the longer of which is associated with proliferating cancer cells (Chierico et al., 2017). *PRMT1* knockdown in LNCaP cells promotes exclusion of *MKI67* exon 7, leading to increased expression of the short isoform (*MKI67-S*) and decreased expression of the pro-proliferative long isoform (*MKI67-L)* (Figures S3C and S3D). Altogether, these data indicate that PRMT1 critically regulates the expression and splicing patterns of AR target genes in prostate cancer cells.

### AR genomic occupancy is impaired by PRMT1 inhibition

Given our observation that t-ARBSs are enriched in proximity to genes downregulated by *PRMT1* knockdown, we next mapped the AR cistrome upon genetic or pharmacologic PRMT1 inhibition. We performed AR chromatin immunoprecipitation and sequencing (ChIP-seq) in LNCaP cells expressing either a control shRNA (sh*LacZ*) or an shRNA targeting PRMT1 (sh*PRMT1*). We observed 33,419 AR peaks in sh*LacZ* infected cells, of which nearly half (16,151, 48%) were lost in sh*PRMT1* infected cells (Figure 4A). Control and *PRMT1* knockdown conditions shared 17,268 peaks (52% of sh*LacZ*; 92% of sh*PRMT1*), while 1,440 peaks (8% of sh*PRMT1*) were gained by *PRMT1* knockdown. These proportions were recapitulated with chemical inhibition of PRMT1 using furamidine (Figure 4B). Furthermore, we observed significant overlap between AR peaks lost with either genetic or pharmacologic inhibition of PRMT1. Of 28,255 AR peaks shared between LNCaP/sh*LacZ* and LNCaP/DMSO experiments, 12,387 (44%) were lost with *PRMT1* knockdown and 8,028 (28%) were lost with furamidine treatment, with an overlap of 5,485 lost peaks between the two conditions (*P* ~ 0 by hypergeometric test). We also observed a decrease in AR binding density across both lost and shared peaks using either chemical or genetic PRMT1 inhibition (Figures 4C and 4D). Next, we assessed whether PRMT1 inhibition leads to global attenuation of AR binding or whether it reprograms the AR cistrome by retargeting AR to alternative target sites. We analyzed the genomic distribution and motif preference of AR target sites in each condition and found that both the genomic locations of AR binding sites and the top enriched AR-bound motifs were largely unaffected by genetic or pharmacologic PRMT1 inhibition (Figures S4A-S4D). We conclude that PRMT1 inhibition impairs AR binding to its canonical target sites but does not dramatically affect AR binding sequence or site preferences.

**Figure 4.**
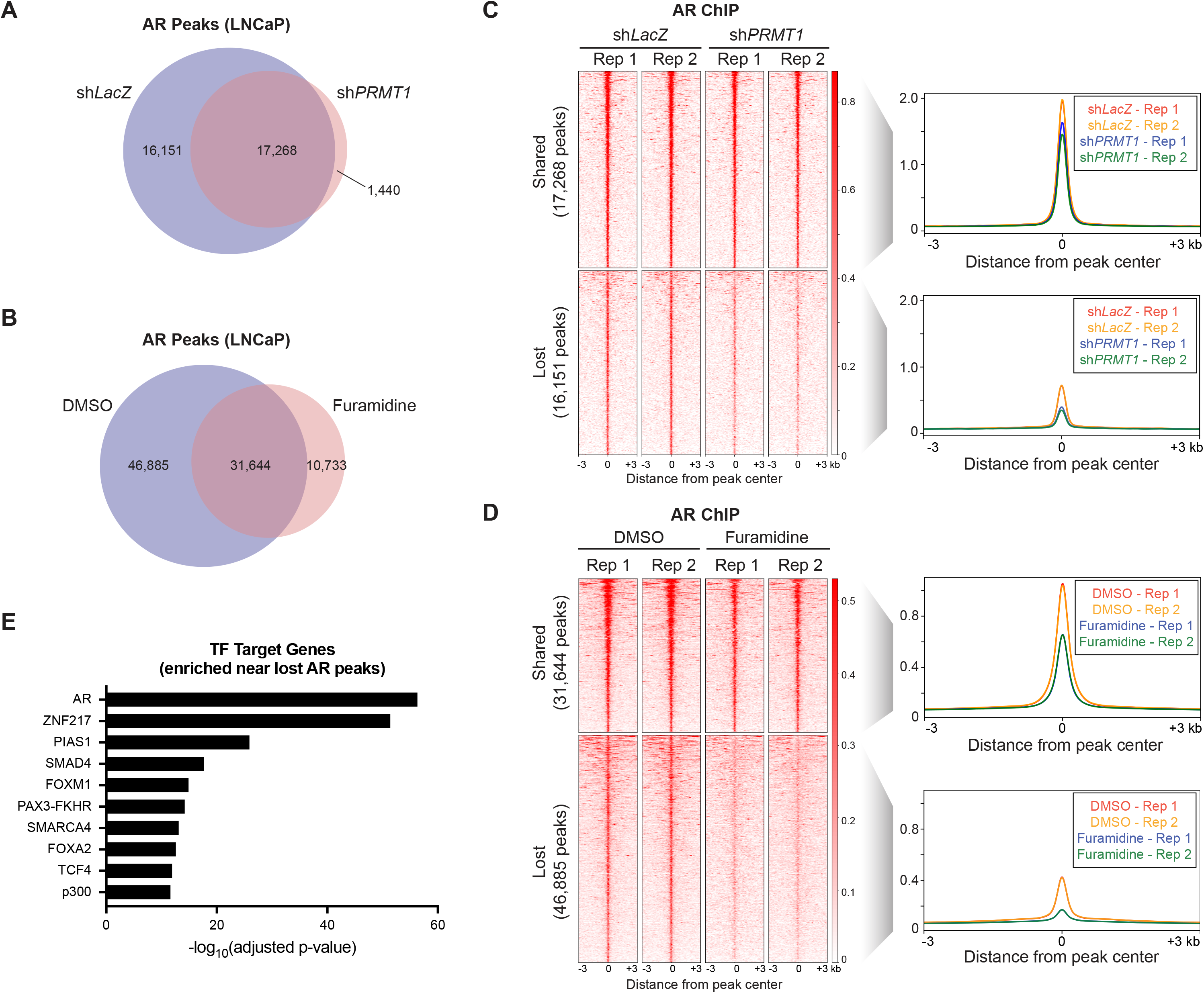
Inhibition of PRMT1 impairs AR binding to genomic target sites. (A and B) Venn diagram showing overlap of AR binding sites identified by AR ChIP-seq in LNCaP cells expressing either control shRNA (sh*LacZ*) or sh*PRMT1* (A) or LNCaP cells treated with either DMSO or furamidine (8 μM) (B). (C) Heatmap of AR binding density over AR peaks shared between sh*LacZ* and sh*PRMT1* conditions (top) or lost upon *PRMT1* knockdown (bottom). Peaks are rank-ordered by AR signal within 3 kb flanking the peak center. Profile plots on the right show average AR ChIP-seq signal in the regions displayed in the heatmaps. Two replicates are shown for each condition. (D) Same data representation as in (C) for AR peaks shared between DMSO and furamidine treatment (top) or lost upon furamidine treatment (bottom). (E) Enrichment analysis of genes located in proximity to AR peaks that were lost by both furamidine treatment and *PRMT1* knockdown. Top enriched transcription factor target gene sets are shown.

Consistent with a prior report (Yu et al., 2010), we noted that the distribution of AR target sites favored distal intergenic regions—where many regulatory enhancer elements reside—over promoter proximal regions (Figures S4A and S4B). Using ChIP-qPCR, we validated that AR binding was selectively enriched at known enhancer elements upstream of *AR* and *KLK3* compared to canonical promoter target sites of AR (Figures S4E and S4F). We also observed diminished AR binding to these regions upon *PRMT1* knockdown or furamidine treatment. Interestingly, we noted that AR ChIP-seq peaks lost by both genetic and pharmacologic targeting of PRMT1 were also enriched in proximity to SMARCA4 and p300 target genes (Figure 4E and Table S5). SMARCA4 is a component of mammalian SWI/SNF complexes and p300 is a histone acetyltransferase. Both participate in chromatin remodeling and modulate the accessibility—and consequently the activity—of regulatory enhancer elements. Furthermore, both have been reported to interact with AR transcriptional complexes and regulate transactivation of AR target genes (Huang et al., 2003). Indeed, we observed significant overlap between AR peaks obtained in our study and both p300 and SMARCA4 peaks previously reported in LNCaP cells. For example, 1,011 of 4,290 (24%) p300 peaks from a prior study (Wang et al., 2011) overlapped with AR peaks shared between the LNCaP/sh*LacZ* and LNCaP/DMSO conditions in our study. Similarly, 6,374 of 7,409 (86%) SMARCA4 peaks from a prior study (Stelloo et al., 2018) overlapped with AR peaks in our study (*P* ~ 0 for both overlaps by hypergeometric test). Together, these data suggest that PRMT1 plays a role in the modulation of AR activity at enhancers, perhaps in concert with other known regulators of this activity.

### *PRMT1* suppression leads to loss of AR genomic occupancy at lineage-specific enhancers and decreased AR target gene expression

We hypothesized that in the context of *PRMT1* suppression, impaired AR binding at enhancer elements may result in decreased enhancer activity and reduced expression of critical oncogenes. The presence of H3K27 acetylation (H3K27ac) in distal enhancer regions can be used to distinguish active from poised enhancers (Creyghton et al., 2010; Rada-Iglesias et al., 2011). We therefore performed H3K27ac ChIP-seq in LNCaP cells to evaluate how PRMT1 regulates the activity of AR target enhancers. Integration of our AR and H3K27ac ChIP-seq data revealed that roughly half of all AR peaks overlapped with H3K27ac peaks in control shRNA-treated LNCaP cells (17,211/31,809 peaks, 54%). Of these, 6,788 peaks (39%) were lost upon *PRMT1* knockdown (Figure 5A). Similar patterns were observed using chemical inhibition of PRMT1 with furamidine (Figure S5A). As with AR binding sites, there was significant overlap between H3K27ac sites lost with *PRMT1* knockdown and furamidine treatment. Of 50,439 H3K27ac peaks shared between LNCaP/sh*LacZ* and LNCaP/DMSO experiments, 6,814 were lost with sh*PRMT1* and 10,250 were lost with furamidine treatment, with an overlap of 3,757 lost peaks between the two conditions (*P* ~ 0 by hypergeometric test).

**Figure 5.**
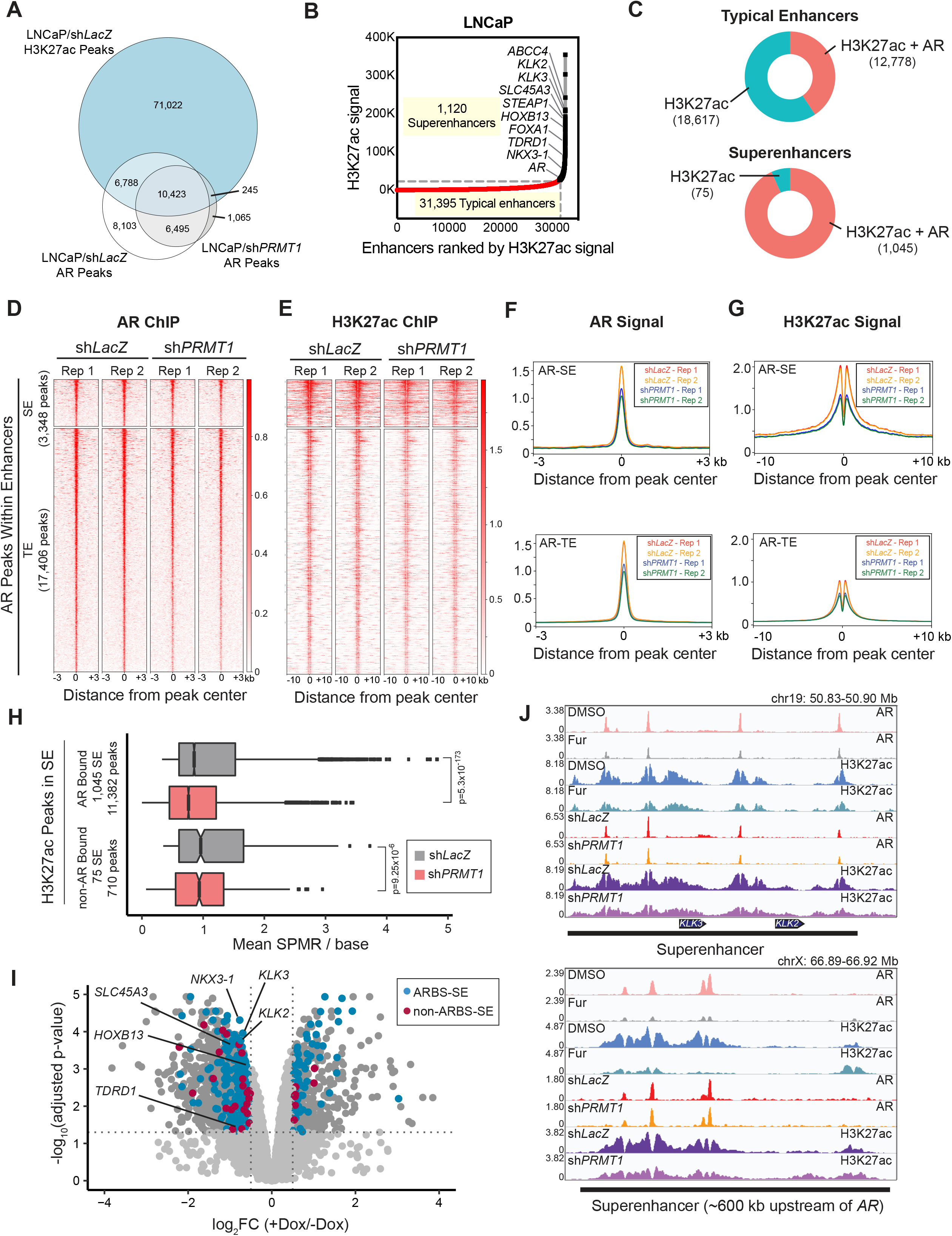
Suppression of *PRMT1* decreases AR target gene expression through reduced AR occupancy and H3K27 acetylation at lineage-specific enhancers. (A) Venn diagram showing the overlap of H3K27ac peaks in LNCaP cells with AR peaks in LNCaP cells expressing sh*LacZ* or sh*PRMT1*. (B) Distribution of H3K27ac ChIP-seq signal across 32,515 typical enhancers (TEs) or superenhancers (SEs) in LNCaP cells. The 1,120 SEs, characterized by high H3K27ac signal, are colored in black. Gene labels indicate SEs proximal to *AR* or AR target gene loci. (C) Donut plots showing the proportion of TEs (top) or SEs (bottom) occupied by AR. (D and E) Heatmaps of AR (D) and H3K27ac (E) ChIP-seq signal over AR peaks in SE or TE regions, shown in the context of either control shRNA or sh*PRMT1*. Peaks are rank-ordered by AR signal within 3 kb of the peak center. H3K27ac signal is shown within 10 kb flanking the peak center. Two replicates are shown for each condition. (F and G) Profile plots of average AR (F) or H3K27ac (G) signal in the regions shown in (D) and (E). (H) Boxplots showing average signal per million reads (SPMR) per base over each H3K27ac peak within AR-occupied or non-AR-occupied SEs in cells expressing sh*LacZ* or *sh*PRMT1. Data represent average of two replicates. p-values were calculated by Mann-Whitney U test. (I) Volcano plot of differentially expressed genes upon *PRMT1* knockdown as determined by transcriptome sequencing. Genes meeting significance and differential expression thresholds of adjusted p-value < 0.05 and |log_2_FC| > 0.5 are colored in dark gray. Of these, genes proximal to AR-occupied SEs (ARBS-SE) are shown in blue while those proximal to non-AR-occupied SEs (non-ARBS-SE) are shown in red. Fisher’s exact test was used to determine enrichment of ARBS-SE-proximal genes (p = 0.029) or non-ARBS-SE-proximal genes (p = 0.131) among those downregulated by *PRMT1* knockdown. Selected canonical *AR* target genes located near SE in LNCaP cells are labeled. (J) AR and H3K27ac ChIP-seq signals at SE regions regulating *KLK2*, *KLK3* (top) or *AR* (bottom) expression are shown in the context of DMSO or furamidine (Fur) treatment, or control or *PRMT1* knockdown. Superenhancers are indicated by black bars. Signals represent average of two replicates.

To further assess AR occupancy in enhancer regions, we first used H3K27ac signal to annotate and rank active enhancers in LNCaP cells. Of the 32,515 active enhancers that were identified, 1,120 were classified as superenhancers (Figure 5B and Table S6). Superenhancers (SEs) are large clusters of enhancers characterized by high transcriptional activity that coordinate the regulation of critical cell identity genes; in cancer cells, SEs are transcriptional hubs that maintain high-level expression of key oncogenic drivers, including lineage-specific oncogenes (Hnisz et al., 2013; Lovén et al., 2013). We noted that *AR* itself and several canonical AR target genes were located in proximity to SE regions in LNCaP cells, and likewise that a large proportion of SEs were occupied by AR (Figures 5B and 5C), in agreement with a prior report (Rasool et al., 2019). Across all AR-occupied enhancers, we observed global decreases in AR and H3K27ac signal upon *PRMT1* knockdown or furamidine treatment (Figures 5D-5G and S5B-S5E). Notably, loss of H3K27ac signal appeared more pronounced at SEs than at typical enhancers (TEs). We also observed a more significant decrease in H3K27ac signal at AR-occupied SEs than at non-AR-occupied SEs upon *PRMT1* knockdown (Figure 5H). As enhancer activity critically affects gene expression, we integrated our ChIP-seq and RNA-seq data to assess whether the observed decreases in AR and H3K27ac signal at SE corresponded to decreases in gene expression. We found that genes proximal to AR-occupied SEs were significantly enriched among those downregulated upon *PRMT1* knockdown; however, genes proximal to non-AR-occupied SEs were not significantly enriched among downregulated genes (Figure 5I), suggesting that PRMT1 regulates the expression of key lineage oncogenes by modulating AR target SE activity. Finally, we evaluated AR and H3K27ac signal at SE regions regulating the expression of *KLK2*, *KLK3*, and *AR* itself, and found decreased AR and H3K27ac signal over these regions in the presence of sh*PRMT1* or furamidine treatment (Figure 5J). We also validated this result by ChIP-qPCR (Figures S5F and S5G; see also Figures S4E and S4F).

PRMT1 has been previously reported to activate gene expression by asymmetric dimethylation of H4R3 (H4R3me2a) (Strahl et al., 2001; Wang, 2001) and by direct association with transcription factor complexes as a transcriptional coactivator (Koh et al., 2001). We therefore sought to test both of these activities of PRMT1 as they might pertain to AR transcriptional complexes in LNCaP cells. *PRMT1* knockdown resulted in globally decreased H4R3me2a (Figure S6A), confirming the importance of PRMT1 for maintenance of this activating transcriptional mark. Global H4R3me2a ChIP-seq profiles have not been previously reported, likely due to a lack of suitable antibodies; we too were unable to obtain reliable H4R3me2a ChIP-seq data (data not shown).

We next sought to assess the interaction between PRMT1 and AR in LNCaP cells. By co-immunoprecipitation, we found that PRMT1 associates with AR on chromatin (Figure S6B), consistent with a prior report that AR and other nuclear hormone receptors interact with PRMT1 *in vitro* (Koh et al., 2001), and suggesting that PRMT1 modulates AR activity through direct interaction with AR transcriptional complexes. Given our prior observation that AR primarily localizes to distal enhancer elements, as well as our observation that PRMT1 regulates AR activity at enhancers, we sought to determine whether PRMT1 might mediate looping between AR-occupied enhancers and target gene promoters; a similar role for PRMT1 has previously been reported at the β-*globin* locus in erythroid progenitor cells (Li et al., 2010). We assessed interaction of the *AR* promoter with its upstream enhancer using chromosome conformation capture (3C) as previously described (Takeda et al., 2018), but did not observe a significant difference in interaction frequency upon *PRMT1* knockdown (Figure S6C).

### *PRMT1* is a selective dependency of *AR*-expressing prostate cancer cell lines

Having established PRMT1 as a critical mediator of AR target gene expression, we next investigated whether this might implicate PRMT1 as a selective vulnerability of AR-driven prostate cancer cells. Consistent with this hypothesis, significant growth inhibition was observed upon *PRMT1* knockdown in *AR*-expressing LNCaP, 22Rv1, and VCaP cells, but not in non-*AR*-expressing PC3 cells (Figures 6A-6C). We also assessed the responses of these cell lines to the PRMT1 inhibitors furamidine and MS023, both of which recapitulate the global loss of PRMT1 catalytic activity observed with *PRMT1* knockdown (Figures S7A and S7B). We observed heightened sensitivity to furamidine or MS023 in *AR*-expressing cells compared to non-*AR*-expressing cells (Figures 6D and S7C). These results suggest that the growth-inhibitory effects of PRMT1 inhibition in prostate cancer cells are specifically mediated through the AR axis.

**Figure 6.**
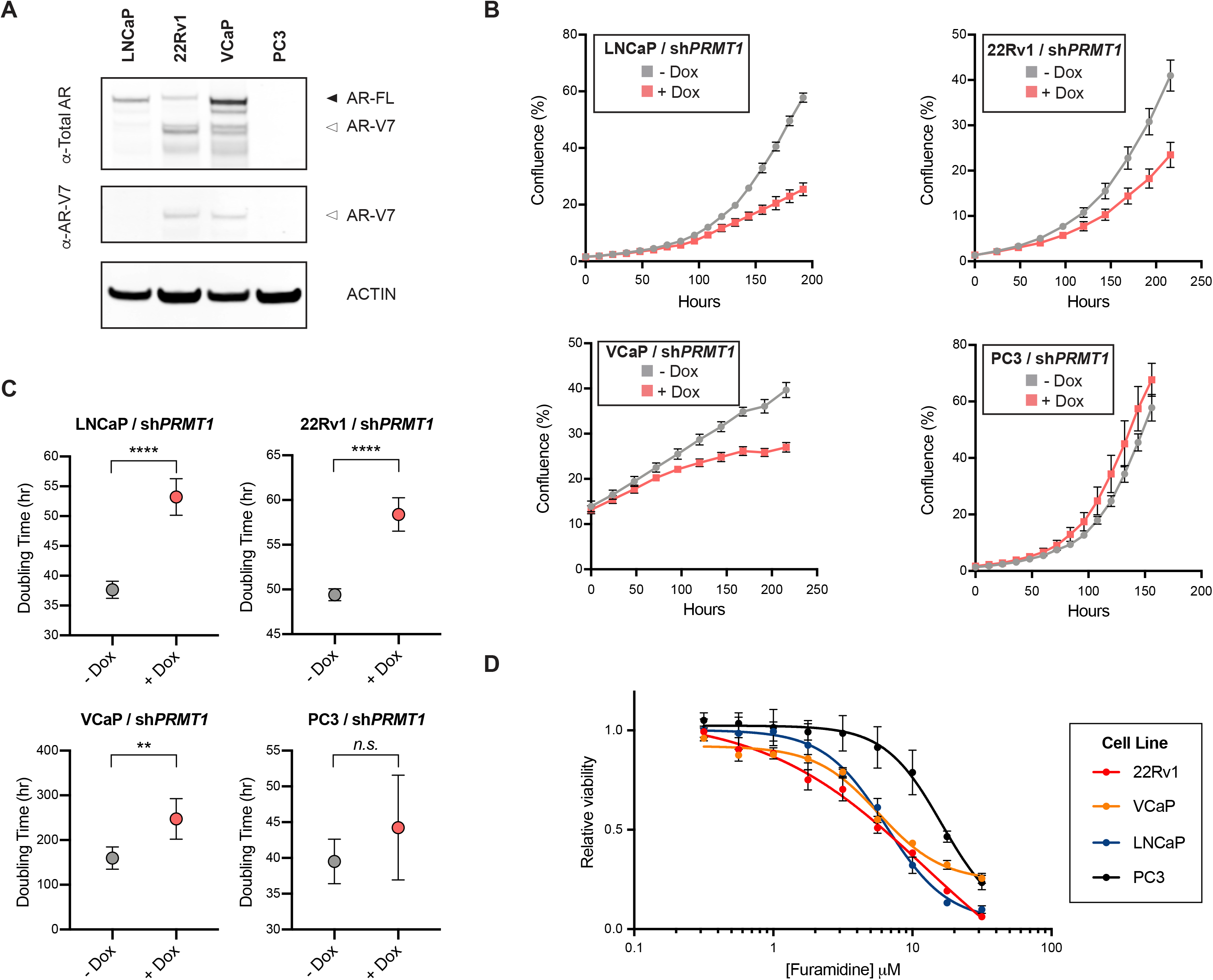
*AR*-expressing prostate cell lines exhibit selective dependency on *PRMT1*. (A) Western blot showing relative *AR-FL* and *AR-V7* expression in parental prostate cancer cell lines. (B) Proliferation of *AR*-expressing (LNCaP, 22Rv1, VCaP) or non-*AR*-expressing (PC3) cell lines with or without dox-induced *PRMT1* knockdown. Confluence readings were taken using an IncuCyte live-cell imager. Error bars represent mean ± SD of the following numbers of replicates: n = 4 (LNCaP), n = 8 (22Rv1), n = 4 (VCaP), n = 3 (PC3). (C) Doubling times of prostate cancer cell lines with or without *PRMT1* knockdown, estimated by nonlinear regression of confluence readings shown in (A). Data are presented as mean with 95% CI. Statistical significance was determined by Student’s t-test. (D) Relative viability (normalized to DMSO) of prostate cancer cell lines after 5 days of treatment with furamidine at the indicated concentrations. Error bars represent mean ± SD, n = 3 replicates. **p < 0.01, ****p < 0.0001.

### Combined AR and PRMT1 targeting leads to synergistic growth inhibition in CRPC cells

Castration-resistant prostate cancer cells commonly exhibit sustained AR signaling despite androgen suppression to castrate levels, which has provided the rationale for development of more potent androgen pathway inhibitors to treat CRPC (Watson et al., 2015). We therefore sought to evaluate whether co-targeting of AR and PRMT1 might suppress the growth of CRPC cells driven by enhanced *AR* or *AR-V* expression. We treated a panel of prostate cancer cell lines (LNCaP, VCaP, 22Rv1, and PC3) with the AR antagonist enzalutamide and the PRMT1 inhibitor furamidine. We also tested this combination in the LNCaP/AR-Enh cell line, an isogenic derivative of parental LNCaP cells in which a second copy of the *AR* enhancer was introduced to the endogenous locus by genetic engineering (Takeda et al., 2018). *AR* enhancer duplication is an exceptionally pervasive somatic alteration in CRPC, found in up to 85% of cases (Dessel et al., 2019; Viswanathan et al., 2018a). We observed synergistic growth inhibition with furamidine and enzalutamide co-treatment in VCaP and 22Rv1 cells, which co-express *AR* and *AR-V*s; notably, 22Rv1 cells are enzalutamide-resistant at baseline owing to high levels of *AR-V7* (Li et al., 2013). In contrast, neither inhibitor substantially affected the viability of PC3 cells, which are AR-negative (Figures 7A and 7B). While enzalutamide alone was sufficient to suppress growth of androgen-sensitive LNCaP cells (which exclusively express *AR-FL*), LNCaP/AR-Enh cells displayed relative resistance to enzalutamide owing to increased *AR* expression, as expected (Takeda et al., 2018). However, we observed re-sensitization of LNCaP/AR-Enh cells to enzalutamide in the context of combined AR and PRMT1 inhibition (Figure 7B), suggesting that blunting of AR signaling via PRMT1 inhibition lowers AR activity below the threshold at which antagonists are again active. Supportive of this notion, we observed a reduction in AR protein in LNCaP/AR-Enh cells upon furamidine treatment, bringing levels to within the range of parental LNCaP cells (Figures 7C and 7D).

**Figure 7.**
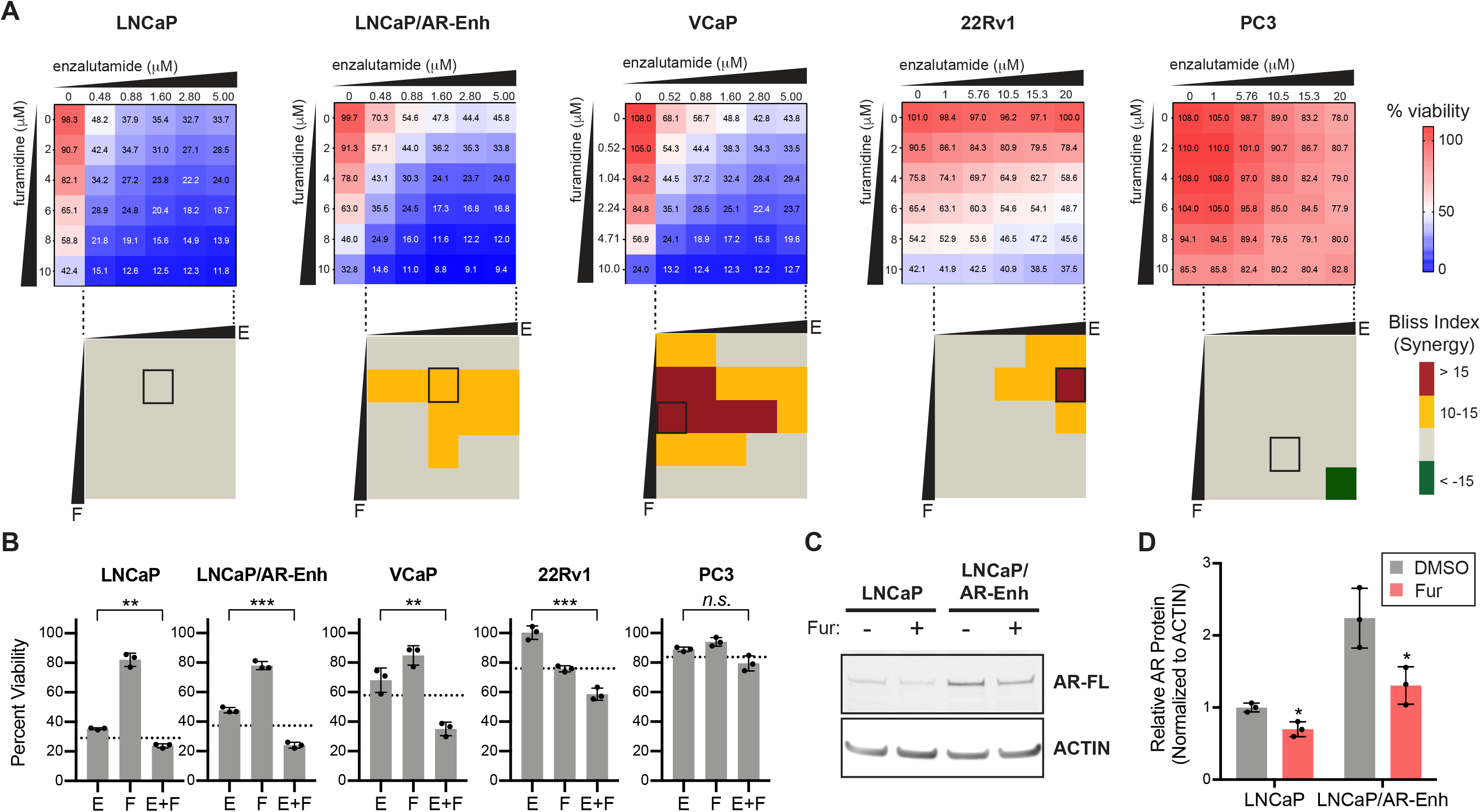
Inhibition of PRMT1 synergizes with enzalutamide to suppress growth of CRPC cells driven by enhanced AR signaling. (A) Heatmaps showing percent viability of prostate cancer cell lines after 7 days of combination treatment with the indicated doses of furamidine and enzalutamide. LNCaP/AR-Enh cells are derived from the parental LNCaP line and contain knock-in of an additional copy of the *AR* enhancer (Takeda et al., 2018). Viabilities are shown relative to the DMSO condition. Quantized heatmaps of Bliss synergy index are shown below each cell line. A box is drawn around the dose combination in each cell line that resulted in the maximum Bliss synergy score. Data represent the mean of n = 3 replicates. (B) Percent viabilities for single-agent compared to combination treatment at the doses indicated by boxes in (A) for each cell line. Dotted lines indicate predicted additive effect of enzalutamide (E) and furamidine (F), calculated by multiplying the percent viabilities upon single-agent treatment at the respective doses. Error bars represent mean ± SD, n = 3 replicates. (C) Western blot showing relative AR protein levels in LNCaP and LNCaP/AR-Enh cells upon furamidine treatment in the context of androgen depletion. Cells were seeded in media supplemented with charcoal stripped serum and treated with DMSO or furamidine (8 μM) for 5 days. (D) Densitometric quantification of western blot in (C). AR protein levels are normalized to actin and shown relative to parental LNCaP treated with DMSO. Error bars represent mean ± SD, n = 3 replicates. Statistical significance was determined by Student’s t-test. *p < 0.05, **p < 0.01, ***p < 0.001, ****p < 0.0001.

## DISCUSSION

Androgen ablation was first shown to be an effective treatment for prostate cancer over eight decades ago (Huggins and Hodges, 1941). Clinical, genomic, and functional studies over the past two decades have converged on a central role for AR in prostate cancer pathogenesis across disease states, including in CRPCs that have developed resistance to primary hormonal therapy (Chen et al., 2004; Takeda et al., 2018; Taplin et al., 1995; Tran et al., 2009; Visakorpi et al., 1995; Viswanathan et al., 2018a). Next-generation androgen pathway inhibitors that inhibit androgen synthesis (e.g. abiraterone) or act as potent AR antagonists (e.g. enzalutamide) are effective for some time in CRPC, but resistance inevitably emerges, most commonly via re-activation of AR signaling. This provides a compelling rationale for the development of orthogonal strategies to target AR output in prostate cancer. Here, we leverage genome-scale genetic screening to systematically identify regulators of *AR* expression, and uncover PRMT1 as a critical component of the AR axis.

Re-activation of AR signaling in prostate cancer may occur through various genetic and non-genetic mechanisms, all of which serve to increase AR levels and/or activity and enable sustained signaling despite low levels of circulating androgen ligands. The most pervasive mechanisms of AR re-activation include copy number amplification of the *AR* gene (Visakorpi et al., 1995) and/or its enhancer (Takeda et al., 2018; Viswanathan et al., 2018a), or the production of *AR* splice variants lacking the C-terminal ligand binding domain of full-length AR (Antonarakis et al., 2014; Dehm et al., 2008). *AR-V*s may be produced either by genomic rearrangements at the *AR* locus (Li et al., 2020) or by aberrant regulation of transcription or splicing (Liu et al., 2014). While the most well-studied truncated AR variant has historically been *AR-V7*, emerging data indicate that prostate cancer cells may express multiple truncated *AR* variants, and that one or more may act coordinately to promote ligand-independent signaling (Kallio et al., 2018; Xu et al., 2015). Moreover, *AR-V7* production in the context of ADT has been shown to be coupled to transcription initiation and elongation rates, indicating that factors that control transcription of the *AR* gene can indirectly control the expression of its splice variants by modulating splicing factor recruitment to the pre-mRNA (Liu et al., 2014). Importantly, the CRISPR/Cas9 screen performed in our study captures control of *AR-V7* expression at both the transcriptional and post-transcriptional levels. Using this approach, we identified factors that coordinately regulated both *AR-FL* and *AR-V7* as well as factors that exhibited selective regulation of *AR-V7*. The former represent the most attractive therapeutic targets, as they may control the expression of not only *AR-V7* but also other *AR* splice isoforms (including *AR-FL*). However, the latter also deserve further study and may elicit important insights into mechanisms of aberrant *AR* splicing in advanced prostate cancer.

In this study, we identified and characterized PRMT1 as a key regulator of AR signaling in CRPC. We show that PRMT1 regulates *AR/AR-V7* expression, as well as AR output more broadly, by influencing the activity of AR at its target enhancers; key among these is the *AR* enhancer itself. At this juncture, the critical substrates of PRMT1 that mediate AR signaling remain unknown. PRMT1 has a multitude of nuclear and non-nuclear substrates, which vary by cell type and context, and one or more of these may be important for the phenotypes observed herein (Hsu et al., 2017). For example, PRMT1 can directly modify histone H4 at the R3 position, and the resulting asymmetric dimethyl mark (H4R3me2a) is thought to facilitate transcriptional activation (Strahl et al., 2001; Wang, 2001). Additionally, PRMT1 is known to modify the chromatin-associated protein CHTOP (one of the top hits in our screen), which may recruit PRMT1 to sites of 5-hydroxymethylcytosine and promote H4R3me2a deposition (Dijk et al., 2010). We also show here that PRMT1 associates with AR on chromatin, consistent with a prior report of PRMT1 as a coactivator of nuclear hormone receptors (Koh et al., 2001). Notably, PRMT1 has also been reported to associate with p160 coactivator proteins, which facilitate assembly of AR transcriptional complexes and bridging of AR-bound promoter and enhancer elements (Koh et al., 2001; Shang et al., 2002). This circumstantial evidence, together with the observation that *AR*-expressing prostate cancer cells exhibit selective dependency on PRMT1, suggests that PRMT1 may modify key components of the AR transcription factor complex. Potential substrates include AR coregulators such as FOXA1 and HOXB13, or perhaps AR itself. Interestingly, recent studies have reported symmetric dimethylation of AR by PRMT5 (Mounir et al., 2016) and asymmetric dimethylation of the progesterone receptor, another nuclear hormone receptor, by PRMT1 (Malbeteau et al., 2020). Thus, PRMT1 may regulate AR output both through its direct effects on *AR* expression and through interactions with AR transcriptional complexes. Identification and characterization of AR-dependent PRMT1 substrates is a ripe area for future investigation.

Our study demonstrates that either genetic or pharmacologic inhibition of PRMT1 globally impairs AR occupancy at a majority of its target sites, leading to a loss of H3K27ac at lineage-specific enhancers and reduced expression of critical oncogenes, including *AR* itself. Blocking AR binding to its target sites via inhibition of PRMT1 represents a promising orthogonal approach to target AR signaling. Current strategies for inhibiting AR transcriptional activity in CRPC rely on androgen synthesis inhibitors or AR antagonists, both of which are ineffective against tumors expressing truncated AR splice variants (Antonarakis et al., 2014). In contrast, we find that *AR*-expressing prostate cancer cells exhibit similar sensitivity to PRMT1 inhibition regardless of *AR-V* expression. This finding suggests that truncated AR variants may also depend on PRMT1 for binding to genomic target sites, which underscores the need for further evaluation of PRMT1 as a therapeutic target in CRPC.

PRMT1 is the primary type I PRMT, accounting for up to 90% of ADMA in cells (Tang et al., 2000), though substrate redundancy among PRMTs has been reported (Dhar et al., 2013). Given the large number of PRMT substrates in the cell (Hsu et al., 2017), some of which may be cell-essential, toxicity may be limiting with PRMT1 monotherapy. Combination therapy can yield increased durability of response with an expanded therapeutic window, as evidenced, for example, by the success of CDK4/6 inhibitor-endocrine therapy combinations in breast cancer (Finn et al., 2016). Our data provide initial support for a similar principle in CRPC, leveraging the combination of direct AR inhibition and PRMT1 inhibition.

While furamidine, the PRMT1 tool compound inhibitor used in this study, exhibits at least 15-fold selectivity for PRMT1 over other PRMTs (Yan et al., 2014), other non-PRMT targets of furamidine have been reported (Jenquin et al., 2018). We therefore cannot exclude the possibility that furamidine treatment may induce phenotypic effects due to its activity on targets other than PRMT1. Still, the strong concordance between the results obtained with furamidine treatment and genetic silencing of *PRMT1* suggests that PRMT1 is primarily responsible for the phenotype described herein. The development of potent and specific clinical-grade PRMT1 inhibitors is therefore an active area of research. New PRMT inhibitors are currently being investigated in clinical studies, including a type I PRMT inhibitor in clinical development (Fedoriw et al., 2019; Fong et al., 2019).

Mediators of AR-regulated gene expression represent attractive intervention points in CRPC; while several factors have been reported to enhance AR transcriptional activity, not all are easily amenable to therapeutic targeting by small molecules (Groner et al., 2016; Gui et al., 2019; Hwang et al., 2019; Lee et al., 2019). In addition, we have recently identified *AR* enhancer alterations in up to 85% of CRPC tumors (Viswanathan et al., 2018a). These non-coding alterations, which may be associated with resistance to next-generation anti-androgens (Takeda et al., 2018), have thus far remained untargetable. Here, we demonstrate that PRMT1 inhibition leads to reduced transcriptional activity at lineage-specific enhancers, including the *AR* enhancer. We furthermore show that dual inhibition of AR and PRMT1 is selectively effective in prostate cancer cells bearing *AR* enhancer amplification as compared with isogenic control cells without enhancer amplification. Altogether, our study establishes a preclinical rationale for the development of strategies to coordinately target AR and PRMT1 in the treatment of AR-driven CRPC.

## ACKNOWLEDGEMENTS

We warmly thank Matthew Meyerson for helpful discussions and guidance. We are grateful for assistance from the DFCI Molecular Biology Core Facility, the Broad Institute and Dana Farber Cancer Institute Flow Cytometry Core Facilities and the Broad Institute Genetic Perturbation Platform. This work was supported by the Department of Defense Prostate Cancer Research Program (W81XWH-17-1-0358 to S.R.V.), Prostate Cancer Foundation Young Investigator Award (to S.R.V.), Prostate Cancer Foundation Challenge Award (to M.L.F), American Cancer Society – AstraZeneca (PF-16-142-01-TBE to J.H.H.), and National Institutes of Health/National Cancer Institute (R00 CA208028 to P.S.C.; R01 CA193910 to M.L.F.; R01 CA204954 to M.L.F.; U01 CA176058 to W.C.H.).

## AUTHOR CONTRIBUTIONS

Designed research: S.T., N.Y.M., M.F.N., P.S.C., S.R.V.

Performed research: S.T., N.Y.M., M.F.N., M.K.G., S.A.A., J-H.S., J.H.H., P.S.C., S.R.V.

Contributed new reagents or analytic tools: C.A.S., D.Y.T.

Provided expertise, designed and/or supervised analysis, and edited the paper: J.H.H., S.C.B., J.L., S.A., X.Z., J.G.D., W.C.H., D.Y.T., M.L.F., P.S.C., S.R.V.

Analyzed data: S.T., N.Y.M., M.F.N., J.G.D., P.S.C., S.R.V.

Wrote the manuscript: S.T., P.S.C., S.R.V.

## DECLARATION OF INTERESTS

J.G.D. consults for Tango Therapeutics, Maze Therapeutics, Foghorn Therapeutics, and Pfizer. W.C.H. is a consultant for ThermoFisher, Solvasta Ventures, MPM Capital, KSQ Therapeutics, iTeos, Tyra Biosciences, Jubilant Therapeutics, Frontier Medicine and Parexel.

## METHODS

### Cell Lines

Prostate cancer cell lines were obtained from the American Type Culture Collection and grown in RPMI (LNCaP, 22Rv1, PC3) or DMEM (VCaP) supplemented with 10% FBS, 100 U mL^−1^ penicillin, 100 μg mL^−1^ streptomycin, 2 mM L-glutamine, and 100 μg mL^−1^ Normocin (Invivogen). For experiments involving doxycycline-inducible shRNA expression, Tet System Approved FBS (Takara, #631101) was used. For experiments performed in androgen depleted conditions, RPMI without phenol red (Gibco, #11835030) supplemented with Charcoal Stripped FBS (Sigma-Aldrich, #F6765) was used. Cell lines were authenticated by short tandem repeat profiling and tested periodically for the presence of mycoplasma. LNCaP *AR* enhancer knock-in line (LNCaP/AR-Enh) is a derivative of the LNCaP parental line and has been previously described (Takeda et al., 2018).

### Compounds

All compounds used were obtained from commercial sources and dissolved in DMSO. Enzalutamide was obtained from Selleck (#S1250) and dissolved to a stock concentration of 10 mM. Furamidine was obtained from Tocris (#5202) and dissolved to a stock concentration of 10 mM. MS023 was obtained from Tocris (#5713) and dissolved to a stock concentration of 1 mM.

### Plasmids

For AR-V7-GFP knock-in experiments, Gibson Assembly was used to construct a donor plasmid consisting of a P2A-Zeo-T2A-EGFP cassette flanked by ~1.5 kilobase homology arms for insertion directly upstream of the *AR CE3* stop codon (pHDR-AR-V7-Zeo-EGFP). Immediately downstream of the 3’ homology arm, the plasmid also contains an EFS-BFP expression cassette to counter-select for random integration events. For constitutive shRNA knockdown experiments, shRNAs were cloned into the pLKO.5 vector (Broad Institute, puromycin resistance) with an shRNA targeting *LacZ* used as a negative control. For inducible knockdown experiments, shRNAs were cloned into a Gateway-compatible lentiviral vector (G418 resistance) under the control of a tetracycline-responsive cytomegalovirus promoter as previously described (Viswanathan et al., 2018b). For CRISPR/Cas9 knockout experiments, Cas9 was cloned into a lentiviral vector (blasticidin resistance) under the control of a tetracyline-responsive cytomegalovirus promoter (Viswanathan et al., 2018b). sgRNAs were cloned into lentiGuide-Puro (Addgene, #52963). Target sequences for shRNAs and sgRNAs are listed in Table S7.

### RT-qPCR

RNA was isolated from cells using the RNeasy Plus Mini Kit (QIAGEN, #74136). cDNA was synthesized from 1 μg of total RNA using SuperScript IV VILO Master Mix (Thermo Fisher Scientific, #11756050). 20 ng of cDNA was used as template in qPCR reactions using Power SYBR Green PCR Master Mix (Applied Biosystems, #4367659) and primers as listed in Table S8. qPCR assays were performed on either QuantStudio 6 Flex Real-Time PCR System (Applied Biosystems) or CFX384 Touch Real-Time PCR Detection System (BIO-RAD), according to the manufacturer’s recommended protocol. Relative gene expression was quantitated using the ΔΔCt method with internal normalization against either *GAPDH* or *β-actin*.

### Drug Treatment Assays

Cells were seeded in 96-well plates at 2,000-10,000 cells per well, depending on the cell line. For single-agent dose response assays, furamidine (Tocris, #5202) or MS023 (Tocris, #5713) was added at the indicated concentrations using a D300e Digital Dispenser (Tecan) or by manual serial dilution, with DMSO treatment as a negative control. After 7 days, cell viability was measured with the CellTiter-Glo Luminescent Cell Viability Assay (Promega, #G7571) and normalized to DMSO control wells. For experiments evaluating combined AR and PRMT1 inhibition, cells were treated with a combination of furamidine (Tocris) and enzalutamide (Selleck) doses as indicated. After 7 days, cell viability was measured by CellTiter-Glo and normalized to DMSO wells. Drug synergy was evaluated with a Bliss independence model using Combenefit v2.021 (Di Veroli et al., 2016).

### Cell Proliferation Assays

Cells were seeded in 96-well plates at densities ranging from 1,000-10,000 cells per well, depending on the cell line. For inducible shRNA experiments, the indicated cell lines were transduced with lentivirus encoding doxycycline-inducible shRNA and selected on G418 prior to seeding at equal densities with or without the addition of 100 ng mL^−1^ doxycycline. Cell proliferation was measured using automated confluence readings from an IncuCyte S3 Live-Cell Analysis System (Essen BioScience).

### Reporter Assays

For AR reporter assays, LNCaP cells were transduced with a lentiviral vector expressing Firefly luciferase under the control of an AR-responsive murine mammary tumor virus (MMTV) promoter. For *PRMT1* knockdown experiments, cells were then transduced with a lentivirally-encoded doxycycline-inducible shRNA targeting *PRMT1*, selected on G418, and seeded at equal densities in 96-well plates with or without the addition of 100 ng mL^−1^ doxycycline to the cell culture medium. For PRMT1 inhibition experiments, cells were treated 24 hours after seeding with either DMSO or the indicated concentrations of furamidine. At 7 days after doxycycline induction or 5 days after drug treatment, reporter activity was assessed using the Bright-Glo Luciferase Assay System (Promega, #E2620). Cells were seeded in parallel for viability measurement using the CellTiter-Glo Luminescent Cell Viability Assay (Promega, #G7571) to normalize for differences in cell proliferation between conditions.

### Lentiviral Infection

Lentivirus was produced using HEK293T cells as previously described (Viswanathan et al., 2018b). For lentiviral transduction, lentivirus was added to culture medium together with 8 μg mL^−1^ polybrene (Santa Cruz Biotechnology, #sc-134220) and cells were spin-infected for 30 minutes at 1000 × *g*. Antibiotic selection was started 24 hours after infection.

### Generation of 22Rv1/AR-V7-GFP Knock-in Lines

To generate knock-in lines, 22Rv1 cells were transfected with pX335 (expressing either a single sgRNA or a pair of sgRNAs targeting *AR CE3*) and the pHDR-AR-V7-Zeo-EGFP donor plasmid. Following transfection, cells were selected with 50-400 μg mL^−1^ of zeocin to enrich for correct editing and GFP-positive/BFP-negative cells were single-cell sorted using a Sony SH800 cell sorter. After expansion in culture, individual clones were screened by PCR and flow cytometry to identify clones with precise knock-in of the GFP donor template into *CE3*. Clone 6 (generated by single-nicking with *CE3* sg2) and Clone 9 (generated by double-nicking with *CE3* sg1 and sg2) were used for all further experiments.

### Genome-scale CRISPR/Cas9 Screening

22Rv1 AR-V7 GFP knock-in Clone 6 was first stably transduced with a doxycycline-inducible Cas9 vector (see Plasmids). Cells were then expanded and transduced in biological replicate with lentivirus from the Brunello genome-wide sgRNA library (Doench et al., 2016). Sufficient numbers of cells were infected to achieve a library representation of at least 1,000 cells per sgRNA at a transduction efficiency of approximately 40%. For infections, polybrene was added at 8 μg mL^−1^ and cells were spun for 30 min at 1000 × *g* at 30°C, before being incubated overnight at 37°C. To induce Cas9 expression, doxycycline was added to the media at 100 ng mL^−1^ at the time of infection and maintained throughout the screen. Selection was started 48 hours after infection with 2 μg mL^−1^ puromycin. At two timepoints (5 and 12 days) after infection, cells were harvested and prepared for sorting by staining with 0.5 μg mL^−1^ propidium iodide and passage through a 40 μm mesh filter. Viable PI-negative/GFP-negative and PI-negative/GFP-low populations were isolated with a Sony SH800 cell sorter with gates set using uninfected GFP-positive cells. Genomic DNA was isolated immediately after cell sorting using the QIAamp DNA Blood Mini Kit (QIAGEN, #51104) with yeast RNA (Thermo Fisher Scientific, #AM7118) added as a carrier to improve DNA recovery from low starting cell numbers. PCR amplification of sgRNA sequences from genomic DNA followed by next-generation sequencing was performed as previously described (Doench et al., 2016).

### Flow Cytometry

For arrayed validation of top screen hits, three independent sgRNAs per gene from the Brunello library were individually cloned into lentiGuide-Puro. One negative control non-targeting guide (control98) and one positive control GFP-targeting guide (GFP sg5) were also cloned. 22Rv1/AR-V7-GFP knock-in Clone 6 and Clone 9 cells with inducible Cas9 were then transduced with sgRNA lentivirus. Cells were subsequently selected with 2 μg mL^−1^ of puromycin and Cas9 expression was induced with 100 ng mL^−1^ of doxycycline. At 5 and 12 days after infection, cells were harvested and GFP fluorescence was measured by flow cytometry. For shRNA experiments, Clone 6 and Clone 9 cells were transduced with a lentivirally-encoded doxycycline-inducible shRNA targeting *PRMT1*. After selection on G418, cells were cultured for 9 days with or without shRNA induction using 100 ng mL^−1^ doxycycline. Cells were harvested and fluorescence intensity was analyzed using an LSR Fortessa flow cytometer (BD Biosciences). All flow cytometric data were collected using FACSDiva software v8.0.1 (BD Biosciences) and analyzed using FlowJo software v10.4.2 (FlowJo). Gates for live, single-cell, GFP-negative populations were set using parental 22Rv1 cells as a no-stain control.

### RNA-seq

LNCaP cells transduced with a doxycycline-inducible shRNA targeting *PRMT1* were cultured with or without 100 ng mL^−1^ doxycycline for 7 days. Total RNA was collected from cells using the RNeasy Plus Mini Kit (QIAGEN, #74136) and concentrations were measured using a NanoDrop 8000 Spectrophotometer (Thermo Fisher Scientific). RNA-sequencing libraries were prepared using the KAPA mRNA HyperPrep Kit (Roche) and pooled prior to paired-end 75 bp sequencing on a NextSeq500 (Illumina).

### Chromatin Immunoprecipitation (ChIP)-seq and ChIP-qPCR

For experiments with *PRMT1* knockdown, LNCaP cells were transduced with a lentiviral vector (pLKO.5) encoding an shRNA targeting either *LacZ* or *PRMT1* and selected on puromycin. For experiments with small-molecule PRMT1 inhibition, LNCaP cells were treated with DMSO or furamidine (8 μM). At 7 days after transduction or 5 days after drug treatment, 10 million cells were fixed using 1% formaldehyde (Thermo Fisher Scientific, #BP531-25) for 10 minutes at room temperature followed by quenching with 125 mM glycine (Sigma-Aldrich, #50046). Cells were rinsed twice with PBS and resuspended in 1 mL lysis buffer (1X PBS, 1% NP-40, 0.5% sodium deoxycholate, 0.1% SDS) supplemented with protease inhibitor cocktail (Roche, #11836170001). Chromatin was sheared to 200-500 base pairs using a Covaris E220 sonicator and cleared by centrifugation for 15 minutes at 19,000 × *g*. Antibodies (AR, 9 μg, Abcam, ab74272; H3K27ac, 1 μg, Diagenode, C15410196) were incubated with 40 μL of protein A/G Dynabeads (Thermo Fisher Scientific, #10002D, #10003D) for at least 6 hours at 4°C before overnight incubation at 4°C with sonicated chromatin. Chromatin-bead complexes were washed 5 times with 1 mL LiCl wash buffer (100 mM Tris pH 7.5, 500 mM LiCl, 1% NP-40, 1% sodium deoxycholate) and rinsed twice with 1 mL TE buffer (10 mM Tris pH 7.5, 0.1 mM EDTA). Immunoprecipitated chromatin was resuspended in 100 μL elution buffer (100 mM NaHCO_3_, 1% SDS) and treated with RNase A (Thermo Fisher Scientific, #12091021) for 30 minutes at 37°C. Crosslinks were reversed in the presence of proteinase K (Thermo Fisher Scientific, #25530049) for 16 hours at 65°C, and the eluted DNA was purified using a MinElute PCR Purification Kit (QIAGEN, #28006). Concentrations of ChIP eluates were measured using a Qubit fluorometer (Thermo Fisher Scientific). For ChIP-qPCR, eluates along with their inputs were quantitated by qPCR using primers listed in Table S8. ChIP-seq libraries were prepared using the NEBNext Ultra II DNA Library Preparation Kit (NEB, #E7645). Libraries were analyzed for fragment size using the Bioanalyzer High-Sensitivity DNA Kit (Agilent, #5067-4626) and quantified using the NEBNext Library Quant Kit for Illumina (NEB, #E7630). After pooling, libraries were sequenced on an Illumina NextSeq 500 using single-end 75bp reads.

### Chromosome conformation capture (3C)

LNCaP cells transduced with a doxycycline-inducible shRNA targeting *PRMT1* were grown in the presence or absence of 100 ng mL^−1^ doxycycline for 7 days before harvesting for 3C analysis, which was performed as previously described (Takeda et al., 2018).

### Co-immunoprecipitation

Lysates for co-immunoprecipitation of chromatin-bound proteins were prepared essentially as described above for ChIP. Approximately 20 million cells were fixed in 1% formaldehyde for 10 minutes at room temperature and quenched with 125 mM glycine. After rinsing with ice-cold PBS, cells were resuspended in ChIP lysis buffer and sonicated. Protein A/G Dynabeads (Thermo Fisher Scientific) were incubated for at least 6 hours at 4°C with an antibody against AR (9 μg, Abcam, ab74272) or normal rabbit IgG (Cell Signaling Technology, #2729) before overnight incubation at 4°C with sonicated lysates. Immunoprecipitates were washed 5 times with LiCl wash buffer, rinsed once with PBS, and then resuspended in 1X NuPAGE LDS Sample Buffer (Thermo Fisher Scientific, #NP0008) and boiled for 5 minutes. Eluates along with input were loaded onto Bolt 4-12% Bis-Tris Plus gels (Thermo Fisher Scientific, #NW04120) for SDS-PAGE and Western blotting analysis. Membranes were probed with an antibody against PRMT1 (1:500, Santa Cruz Biotechnology, sc-166963).

### Histone Acid Extraction

LNCaP cells transduced with a doxycycline-inducible shRNA vector targeting *PRMT1* were grown in the presence or absence of 100 ng mL^−1^ doxycycline. After 7 days of shRNA induction, cells were harvested and washed in ice-cold PBS. To extract nuclei, cells were resuspended in Triton Extraction Buffer (1X PBS, 0.5% Triton X-100, 2 mM PMSF, 0.02% NaN_3_) at a density of 10^7^ cells mL^−1^ and incubated with rotation for 10 minutes at 4°C. Nuclei were pelleted by centrifugation at 6,500 × *g* for 10 minutes at 4°C, washed in half the original volume of Triton Extraction Buffer, and pelleted again as before. Nuclei were resuspended in 0.2 M HCl at a density of 4 × 10^7^ nuclei mL^−1^ and histones were acid extracted overnight with rotation at 4°C. Nuclear debris was pelleted by centrifugation at 6,500 × *g* for 10 minutes at 4°C, and the histone-containing supernatant was neutralized using 1/10 volume of 2 M NaOH. Samples were loaded onto Bolt 4-12% Bis-Tris Plus gels (Thermo Fisher Scientific, #NW04120) for SDS-PAGE and Western blotting analysis. Membranes were probed using antibodies against H4R3me2a (1:1,000, Active Motif, #39705) and total H3 (1:1000, Active Motif, #39763).

### Western Blotting

Cells were lysed on ice with RIPA lysis buffer (Thermo Fisher Scientific, #89901) supplemented with protease inhibitors (Roche, #11836170001) and quantitated using a Pierce BCA Protein Assay Kit (Thermo Fisher Scientific, #23225). Equal amounts of protein were loaded onto NuPAGE 4-12% Bis-Tris Protein Gels (Thermo Fisher Scientific, #NP0335) for separation by SDS-PAGE. Proteins were transferred to nitrocellulose membranes using an iBlot2 (Thermo Fisher Scientific), followed by overnight incubation at 4°C with the following primary antibodies and dilutions. AR: Cell Signaling (#5153), 1:2,000. AR-V7: RevMAb (#31-1109-00), 1:500. PRMT1: Cell Signaling (#2449), 1:1,000. ADMA: Cell Signaling (#13522), 1:250. β-Actin: Cell Signaling (#3700), 1:5,000. Membranes were washed in TBS-T before incubation for 1 hour at room temperature with the following secondary antibodies: IRDye 680LT Goat anti-Rabbit IgG: LI-COR (#926-68021), 1:20,000. IRDye 800CW Donkey anti-Mouse IgG: LI-COR (#926-32212), 1:20,000. Immunoblots were imaged with the Odyssey CLx Infrared Imaging System (LI-COR Biosciences). Band intensity was quantitated using ImageStudioLite v5.2.5.

### Pulse-Chase

Gene-specific mRNA synthesis and degradation kinetics were measured by a pulse-chase assay using the Click-iT Nascent RNA Capture Kit (Thermo Fisher Scientific, #C10365) per manufacturer’s instructions. LNCaP cells transduced with a lentivirally-encoded doxycycline-inducible shRNA targeting *PRMT1* were cultured with or without 100 ng mL^−1^ doxycycline. After 7 days of shRNA induction, cells were pulse-labeled with 5-ethynyl uridine (EU) for 4 hours before the label was chased by replacing the growth medium with regular medium not containing EU. RNA was collected from cells using the RNeasy Mini Kit (QIAGEN, #74106) at the indicated time points during the pulse and chase. EU-labeled transcripts were biotinylated in a copper-catalyzed click reaction before isolation with Streptavidin beads according to the manufacturer’s protocol. cDNA was synthesized from bead-captured RNA using SuperScript IV VILO Master Mix (Thermo Fisher Scientific, #11756050) and analyzed by qPCR as described above, using *β-actin* as an internal normalization control.

### Bioinformatic Analyses

#### Calling of hits from genome-scale CRISPR/Cas9 screen

Sequencing data from the CRISPR/Cas9 screen was processed as previously described (Doench et al., 2016). Reads were initially deconvoluted to obtain read counts for each sgRNA in the library. Read counts were then converted to log-norm values by first normalizing to reads per million (RPM) and then log_2_-transforming after adding 1 to each sgRNA to eliminate zero values. The GFP-negative and GFP-low sorted populations were found to have a high degree of agreement in enriched sgRNAs, so to improve hit detection, their read counts were summed prior to log-norm transformation. Enrichment of each sgRNA was then calculated as the log_2_ fold-change relative to its abundance in the original plasmid DNA pool. Finally, replicates for each timepoint were averaged before analysis with STARS v1.2 (Broad Institute) to obtain a ranked gene list. Gene ontology enrichment analysis was performed on all screen hits that scored with FDR < 0.25 at either timepoint using Enrichr (Chen et al., 2013).

#### Clinical Dataset Analysis

For comparison of *PRMT1* mRNA levels between normal tissue, tumor-adjacent normal tissue, primary tumors, and metastatic tumors, expression data for *PRMT1* (microarray probe 60490_r_at) were obtained from published dataset GSE6919 (Chandran et al., 2007). For analysis of disease-free survival following prostatectomy, data were obtained from the TCGA Pan-Cancer Atlas through cBioPortal (Cerami et al., 2012; Gao et al., 2013; Hoadley et al., 2018). mRNA expression z-score thresholds used to define low or high *PRMT1* expression were z < −1 or z > 1, respectively. All cases not meeting these thresholds were defined as having intermediate *PRMT1* expression. For analysis of *AR* and AR target gene expression in castration-resistant prostate cancer tumors with low or high *PRMT1* expression, data were obtained from a published dataset through cBioPortal (Abida et al., 2019). Tumors were classified as ‘*PRMT1* low’ or ‘*PRMT1* high’ using mRNA expression z-score thresholds of z < −1 or z > 1, respectively.

#### RNA-seq Analysis

Paired-end sequencing reads were aligned to the human genome reference build hg38 using STAR v2.7.2 (Dobin et al., 2013). Transcripts were filtered based on read support (sum of read counts across three biological replicates > 30) prior to gene-level and isoform-level differential expression analysis using the voom transformation in limma v3.40.6 (Ritchie et al., 2015). Thresholds for significant down/upregulation were defined as q < 0.05 and |log_2_fold-change| > 0.5. Differentially expressed genes were analyzed for AR target enrichment using a previously described list of genes that are proximal to tumor-specific AR binding sites and overexpressed in tumor compared to normal tissue (Pomerantz et al., 2015). Analysis of differential alternative splicing events was performed using rMATS v4.0.2 (Shen et al., 2014). The rMATS output was filtered to include only events for which the sum of inclusion counts and skipping counts was greater than or equal to 10 for both sets of samples. Significant differential splicing events were defined using q < 0.05, |inclusion level difference| > 0.1. For visualization of alternative splicing, sashimi plots were generated using rmats2sashimiplot v2.0.3. Enrichment analysis was performed on differentially expressed or spliced genes using Enrichr (Chen et al., 2013). For enrichment analysis of genes downregulated upon *PRMT1* knockdown, the top 100 downregulated genes, ranked by t-statistic, were assessed for enrichment of transcription factor target genes based on gene lists from ENCODE and ChEA (Lachmann et al., 2010), or for enrichment of transcription factor co-expressed genes based on expression data from ARCHS4 (Lachmann et al., 2018). For enrichment analysis of genes with significant exon skipping events, the top 250 transcripts ranked by −log_10_(q-value) * ILD were compared to target gene lists from transcription factor ChIP-seq studies in ChEA.

#### ChIP-seq Analysis

ChIP-seq data were processed with the ChiLin pipeline (Qin et al., 2016) in simple mode using bwa to align to the hg38 human reference genome and MACS2 to call peaks, using the ‘narrow’ setting. For each condition, two replicate IP samples were processed along with their corresponding input DNA controls. Peaks were merged prior to calculation of 2-way overlaps using bedtools v2.29.2 (Quinlan and Hall, 2010) with the -u flag. Regions in which peaks were lost or gained upon furamidine treatment or *PRMT1* knockdown were determined using the -v flag. 3-way peak overlaps were calculated using ChIPpeakAnno v3.18.2 (Zhu et al., 2010). For peaks involved in multiple overlaps, each instance of overlap was counted as an individual peak. Motif analysis was performed using HOMER v4.10 (Heinz et al., 2010) with fragment size set to the size of the region being analyzed. ChIPseeker v1.20.0 (Yu et al., 2015) was used to annotate AR peaks with genomic region as well as the nearest gene based on transcription start site. Enrichment analysis was performed on annotated genes proximal to AR peaks using Enrichr (Chen et al., 2013). Genes in proximity to lost AR peaks common to furamidine treatment and *PRMT1* knockdown were assessed against target gene lists from ChIP-seq studies in ChEA (Lachmann et al., 2010). Bedtools was used as described above to assess overlap between AR peaks and p300 and SMARCA4 peaks from published ChIP-seq datasets (Stelloo et al., 2018; Wang et al., 2011). ROSE (Lovén et al., 2013; Whyte et al., 2013) was used to call enhancers and superenhancers based on H3K27ac signals, as well as to annotate superenhancers with nearby genes. Bedtools was used to intersect AR peaks with enhancer or superenhancer regions as described above. Heatmaps and profile plots for data visualization were generated using deepTools v2.5.7 (Ramírez et al., 2016). AR and H3K27ac ChIP-seq signals were centered by peak summit and ranked by average AR signal within a specified window around summit. Average ChIP-seq signal over bed regions was calculated using bigWigAverageOverBed v2. ChIP-seq signals at selected genomic loci were visualized using IGV (Robinson et al., 2011).

### Statistics

Statistical tests, sample sizes, and the resulting p-values are described in figure legends. Error bars represent SD, SEM, or CI from representative experiments repeated as indicated in figure legends. Student’s t*-*tests were performed in Microsoft Excel using the two-tailed distribution and assuming unequal variance. Other statistical tests were performed using GraphPad Prism 7 or R v3.6.1. Significance was assessed based on a p-value threshold of p < 0.05 or as otherwise described.

## SUPPLEMENTAL FIGURES AND TABLES

**Figure S1. Validation of CRISPR/Cas9 screening reagents and top screen hits.**

**Figure S2. PRMT1 regulates *AR/AR-V7* expression.**

**Figure S3. Transcriptome sequencing reveals splicing changes in AR target genes upon *PRMT1* knockdown.**

**Figure S4. Additional analyses related to AR ChIP-seq in LNCaP cells.**

**Figure S5. Small-molecule PRMT1 inhibition leads to loss of AR binding and H3K27 acetylation at enhancer regions.**

**Figure S6. Role of PRMT1 in modulating AR transcriptional activity.**

**Figure S7. Validation of *PRMT1* knockdown and PRMT1 inhibitor activity.**

**Table S1. Screen hits enriched on day 5, ranked by STARS score.**

**Table S2. Screen hits enriched on day 12, ranked by STARS score.**

**Table S3. Differential gene expression analysis of LNCaP cells with or without *PRMT1* knockdown.** Significantly differentially expressed genes (q < 0.05 and |log_2_fold-change| > 0.5) are listed.

**Table S4. rMATS analysis of differential exon inclusion in LNCaP cells with or without *PRMT1* knockdown.** Positive inclusion level difference (ILD) values indicate exons relatively excluded in the context of *PRMT1* knockdown. Significant differential exon inclusion events (q < 0.05) are listed.

**Table S5. Enrichment analysis of lost AR peaks common to furamidine treatment and *PRMT1* knockdown.** Enriched target gene sets from transcription factor ChIP-seq studies are listed.

**Table S6. ROSE analysis of H3K27ac ChIP-seq data from LNCaP cells.** Typical enhancers and superenhancers called by ROSE analysis of LNCaP cells expressing control shRNA (sh*LacZ*) are listed.

**Table S7. shRNA and sgRNA target sequences.**

**Table S8. Primer sequences for qPCR and genotyping assays.**

