## Supplemental Information for "A genome-scale CRISPR screen reveals PRMT1 as a critical regulator of androgen receptor signaling in prostate cancer"

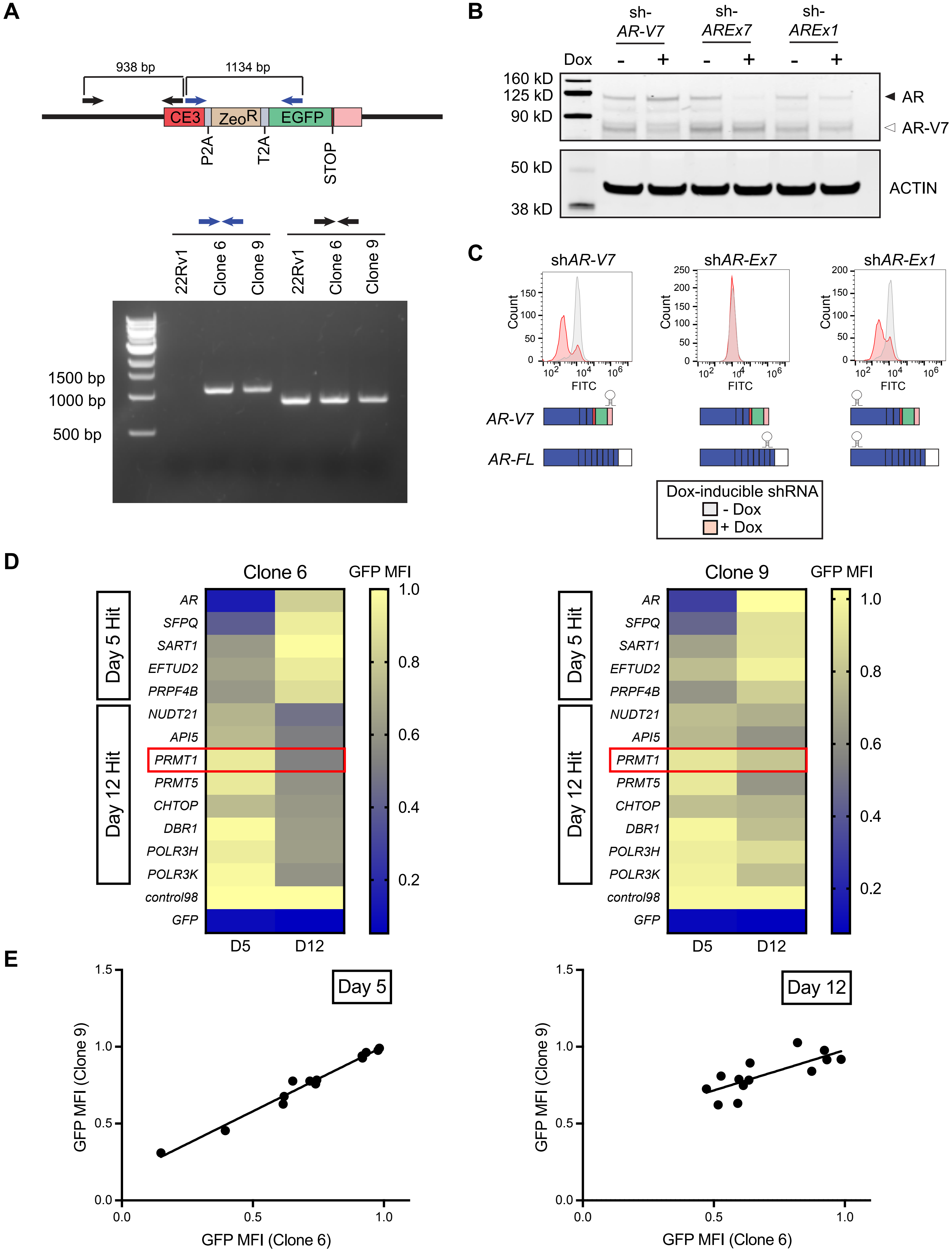

A
938 bp
1134 bp
CE3
ZeoR
EGFP
P2A
T2A
STOP
22Rv1
Clone 6
Clone 9
22Rv1
Clone 6
Clone 9
1500 bp
1000 bp
500 bp
B
sh-AR-V7
sh-AREx7
sh-AREx1
Dox
160 kD
125 kD
90 kD
AR
AR-V7
50 kD
38 kD
ACTIN
C
shAR-V7
shAR-Ex7
shAR-Ex1
Count
FITC
AR-V7
AR-FL
Dox-inducible shRNA
- Dox
+ Dox
D
Clone 6
GFP MFI
Day 5 Hit
Day 12 Hit
AR
SFPQ
SART1
EFTUD2
PRPF4B
NUDT21
API5
PRMT1
PRMT5
CHTOP
DBR1
POLR3H
POLR3K
control98
GFP
D5
D12
Clone 9
E
Day 5
Day 12
GFP MFI (Clone 9)
GFP MFI (Clone 6)

**Figure S1. Validation of CRISPR/Cas9 screening reagents and top screen hits. Related to Figure 1.**
(A) Validation of knock-in integration site in two independent clones of the 22Rv1/AR-V7-GFP reporter line. Integration site was confirmed by PCR on genomic DNA from knock-in clones using genotyping primers positioned as shown, using parental 22Rv1 gDNA as a control.
(B) Validation of shRNA reagents for isoform-specific *AR* knockdown. 22Rv1 cells were transduced with dox-inducible shRNAs targeting *AR-V7* (sh*AR-CE3*), full-length *AR* (sh*AR-Ex7)*, or all *AR* isoforms (sh*AR-Ex1*).
(C) Validation that the knock-in reporter line reports on *AR/AR-V7* expression. AR-V7-GFP levels were measured by flow cytometry after knockdown of *AR* in 22Rv1/AR-V7-GFP cells (Clone 6) using isoform-specific shRNAs as indicated.
(D) Arrayed validation of top hits from genome-scale CRISPR/Cas9 screen by flow cytometry. Heatmap showing median fluorescence intensity (MFI) in two independent clones of 22Rv1/AR-V7-GFP cells at two timepoints after sgRNA-mediated knockout of the indicated screen hits. MFI values are normalized to a negative control sgRNA (*control98*) at each timepoint. Data represent the mean of n = 3 replicates.
(E) Scatterplots showing correlation between GFP MFI in Clone 6 and Clone 9 at day 5 or day 12 after knockout of screen hits.

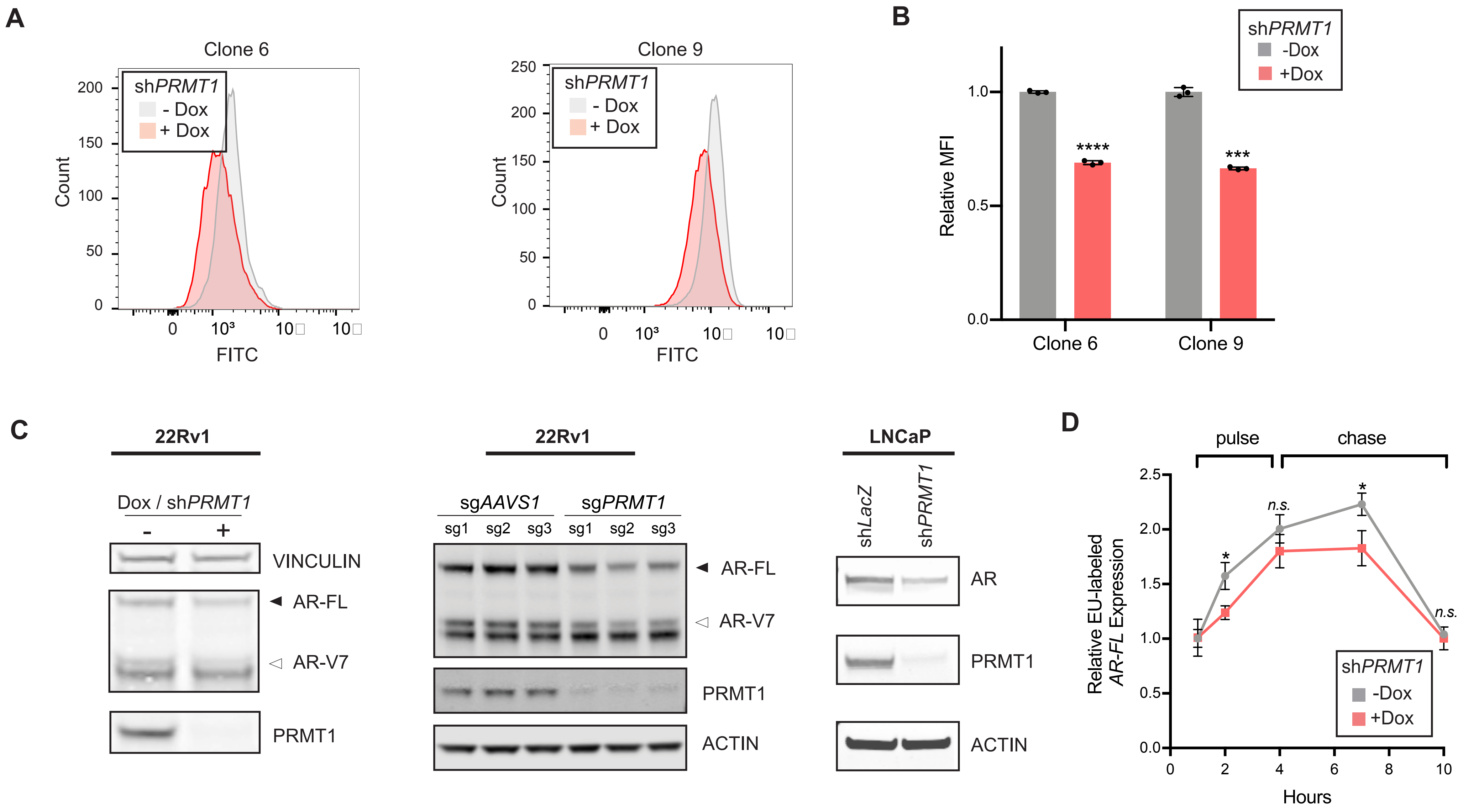


**Figure S2. PRMT1 regulates *AR*/*AR-V7* expression. Related to Figure 2.**
(A) AR-V7-GFP levels in two independent clones of 22Rv1/AR-V7-GFP upon *PRMT1* knockdown using a dox-inducible shRNA. Representative histograms of fluorescence intensity determined by flow cytometry are shown from an experiment conducted in triplicate.
(B) Median fluorescence intensities from the experiment shown in (A), normalized to no dox. Error bars represent mean ± SD, n = 3 replicates.
(C) Validation of decrease in AR-FL and AR-V7 protein in 22Rv1 or LNCaP cells upon *PRMT1* knockdown or knockout. *Left*: Western blot showing decreased AR-FL and AR-V7 protein upon *PRMT1* knockdown in 22Rv1 cells. *Middle*: Western blot showing decreased AR-FL and AR-V7 protein upon control (*AAVS1*) or *PRMT1* knockout in 22Rv1 cells. *Right*: Western blot showing decrease in AR protein in LNCaP cells upon *PRMT1* knockdown.
(D) Ethynyl uridine (EU)-labeled *AR-FL* transcript levels at the indicated timepoints during EU pulse and unlabeled chase of LNCaP cells with or without dox-induced *PRMT1* knockdown. Transcript levels are shown relative to t = 1 hr in each condition. Error bars represent mean ± SD, n = 3 replicates. Statistical significance was determined by Student's t-test. *$p < 0.05$; ***$p < 0.001$; ****$p < 0.0001$.

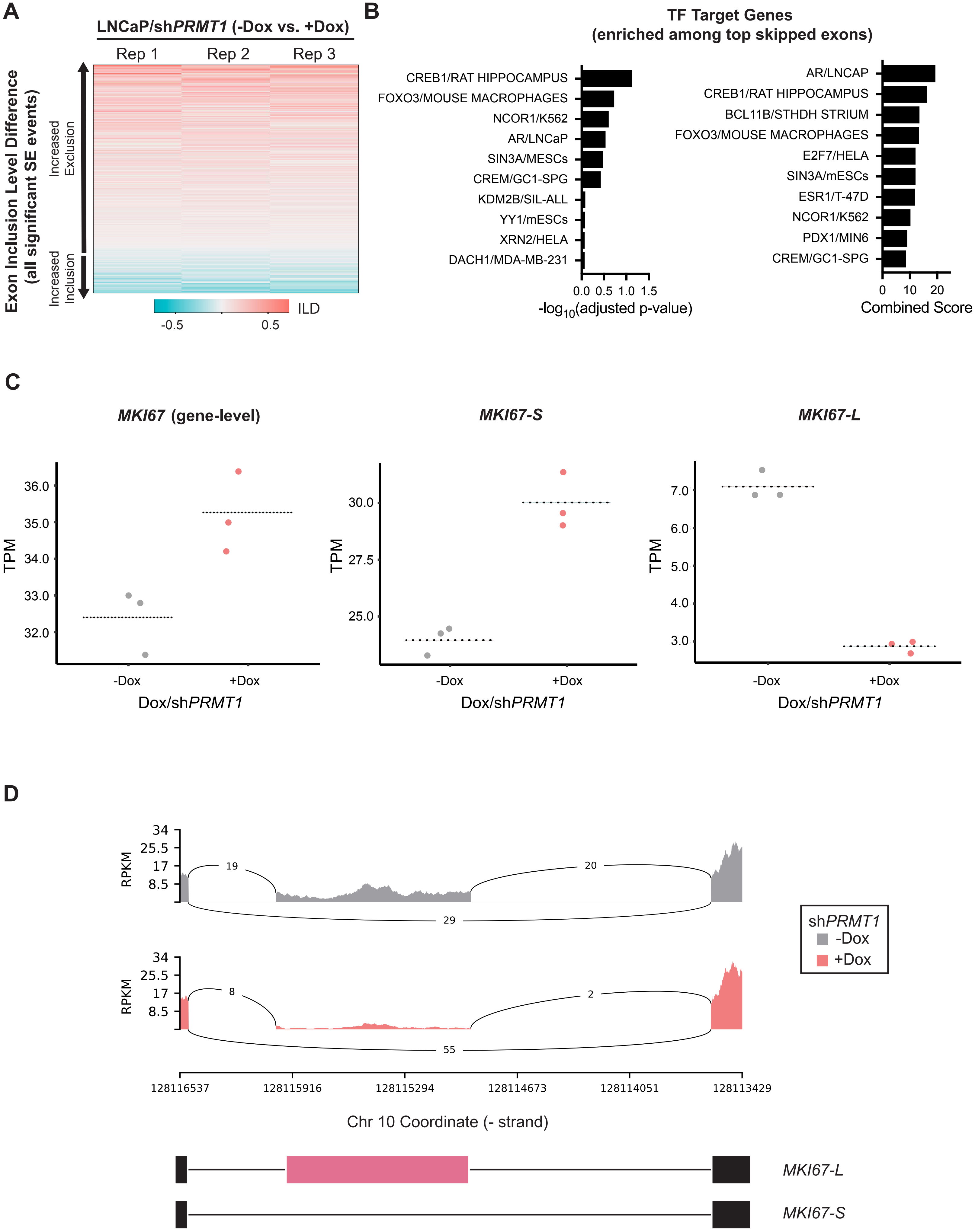

A
LNCaP/shPRMT1 (-Dox vs. +Dox)
Rep 1
Rep 2
Rep 3
Exon Inclusion Level Difference (all significant SE events)
Increased Exclusion
Increased Inclusion
ILD
-0.5
0.5
B
TF Target Genes (enriched among top skipped exons)
CREB1/RAT HIPPOCAMPUS
FOXO3/MOUSE MACROPHAGES
NCOR1/K562
AR/LNCaP
SIN3A/MESCs
CREM/GC1-SPG
KDM2B/SIL-ALL
YY1/mESCs
XRN2/HELA
DACH1/MDA-MB-231
0.0 0.5 1.0 1.5
-log10(adjusted p-value)
AR/LNCAP
CREB1/RAT HIPPOCAMPUS
BCL11B/STHDH STRIUM
FOXO3/MOUSE MACROPHAGES
E2F7/HELA
SIN3A/mESCs
ESR1/T-47D
NCOR1/K562
PDX1/MIN6
CREM/GC1-SPG
0 10 20
Combined Score
C
MKI67 (gene-level)
MKI67-S
MKI67-L
TPM
36.0
35.0
34.0
33.0
32.0
30.0
27.5
25.0
7.0
6.0
5.0
4.0
3.0
-Dox
+Dox
Dox/shPRMT1
D
34
25.5
17
8.5
RPKM
19
20
29
8
2
55
shPRMT1
-Dox
+Dox
128116537
128115916
128115294
128114673
128114051
128113429
Chr 10 Coordinate (- strand)
MKI67-L
MKI67-S

**Figure S3. Transcriptome sequencing reveals splicing changes in AR target genes upon *PRMT1* knockdown. Related to Figure 3.**

(A) Heatmap of inclusion level difference (ILD) for all differentially included exons in LNCaP cells upon *PRMT1* knockdown. Exons are rank-ordered by ILD.

(B) Enrichment analysis of genes containing differentially included exons. Enriched was assessed among target genes from the indicated transcription factor ChIP-seq studies. The top enriched transcription factors ranked by adjusted p-value (left) or combined score (right) are shown. Combined score is based on p-value and z-score of deviation from expected rank.

(C) Expression of the *MKI67* gene and its short (*MKI67-S*) or long (*MKI67-L*) isoforms with or without dox-induced *PRMT1* knockdown. Expression is presented as transcripts per million (TPM). Dotted lines represent the mean of n = 3 replicates in each condition. Expression differences are significant at the gene level (q = 0.030) and isoform level (q = 0.021, *MKI67-S*; q = 0.005, *MKI67-L*).

(D) Sashimi plots around exon 7 of *MKI67* showing differential splicing of *MKI67* pre-mRNA in the presence or absence of *PRMT1* knockdown. Exclusion of exon 7 (pink box in the schematic) generates the short splice isoform (*MKI67-S*). The *y*-axis represents a modified reads per kilobase per million (RPKM) value. Reads shown for each splice event represent the mean of n = 3 replicates.

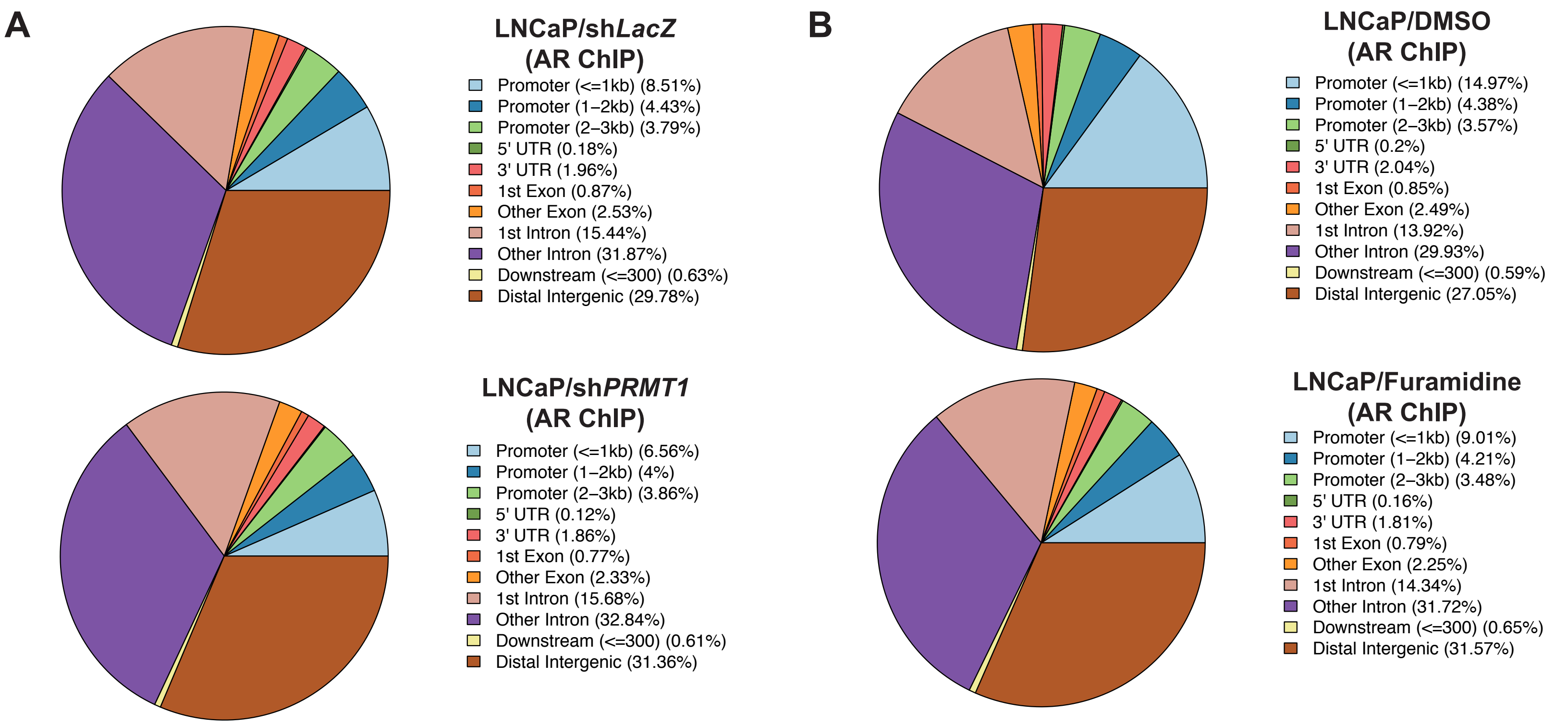


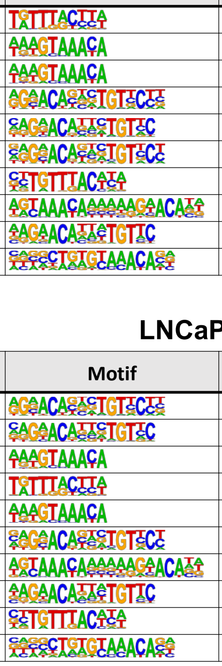

**C**

**LNCaP/shLacZ – AR ChIP**

| Rank | Motif | Name | p-value |
|---|---|---|---|
| 1 | [illegible] | FOXM1 / MCF7-FOXM1-ChIP-Seq (GSE72977) | 1e-2582 |
| 2 | [illegible] | FOXA1 / MCF7-FOXA1-ChIP-Seq (GSE26831) | 1e-2452 |
| 3 | [illegible] | FOXA1 / LNCAP-FOXA1-ChIP-Seq (GSE27824) | 1e-2441 |
| 4 | [illegible] | ARE / LNCAP-AR-ChIP-Seq (GSE27824) | 1e-2327 |
| 5 | [illegible] | GRE / RAW264.7-GRE-ChIP-Seq (Unpublished) | 1e-2327 |
| 6 | [illegible] | GRE / A549-GR-ChIP-Seq (GSE32465) | 1e-2205 |
| 7 | [illegible] | Foxa2 / Liver-Foxa2-ChIP-Seq (GSE25694) | 1e-2070 |
| 8 | [illegible] | FOXA1:AR / LNCAP-AR-ChIP-Seq (GSE27824) | 1e-2041 |
| 9 | [illegible] | PGR / EndoStromal-PGR-ChIP-Seq (GSE69539) | 1e-1948 |
| 10 | [illegible] | Fox:Ebox / Panc1-Foxa2-ChIP-Seq (GSE47459) | 1e-1911 |

**LNCaP/shPRMT1 – AR ChIP**

| Rank | Motif | Name | p-value |
|---|---|---|---|
| 1 | [illegible] | ARE / LNCAP-AR-ChIP-Seq (GSE27824) | 1e-1799 |
| 2 | [illegible] | GRE / RAW264.7-GRE-ChIP-Seq (Unpublished) | 1e-1765 |
| 3 | [illegible] | FOXA1 / MCF7-FOXA1-ChIP-Seq (GSE26831) | 1e-1703 |
| 4 | [illegible] | FOXM1 / MCF7-FOXM1-ChIP-Seq (GSE72977) | 1e-1695 |
| 5 | [illegible] | FOXA1 / LNCAP-FOXA1-ChIP-Seq (GSE27824) | 1e-1670 |
| 6 | [illegible] | GRE / A549-GR-ChIP-Seq (GSE32465) | 1e-1665 |
| 7 | [illegible] | FOXA1:AR / LNCAP-AR-ChIP-Seq (GSE27824) | 1e-1651 |
| 8 | [illegible] | PGR / EndoStromal-PGR-ChIP-Seq (GSE69539) | 1e-1436 |
| 9 | [illegible] | Foxa2 / Liver-Foxa2-ChIP-Seq (GSE25694) | 1e-1351 |
| 10 | [illegible] | Fox:Ebox / Panc1-Foxa2-ChIP-Seq (GSE47459) | 1e-1189 |

**D**

**LNCaP/DMSO – AR ChIP**

| Rank | Motif | Name | p-value |
|---|---|---|---|
| 1 | [illegible] | FOXM1 / MCF7-FOXM1-ChIP-Seq (GSE72977) | 1e-4755 |
| 2 | [illegible] | FOXA1 / MCF7-FOXA1-ChIP-Seq (GSE26831) | 1e-4488 |
| 3 | [illegible] | FOXA1 / LNCAP-FOXA1-ChIP-Seq (GSE27824) | 1e-4366 |
| 4 | [illegible] | Foxa2 / Liver-Foxa2-ChIP-Seq (GSE25694) | 1e-3598 |
| 5 | [illegible] | Fox:Ebox / Panc1-Foxa2-ChIP-Seq (GSE47459) | 1e-3404 |
| 6 | [illegible] | Foxa3 / Liver-Foxa3-ChIP-Seq (GSE77670) | 1e-3205 |
| 7 | [illegible] | FOXA1:AR / LNCAP-AR-ChIP-Seq (GSE27824) | 1e-2228 |
| 8 | [illegible] | FoxL2 / Ovary-FoxL2-ChIP-Seq (GSE60858) | 1e-2046 |
| 9 | [illegible] | FOXK1 / HEK293-FOXK1-ChIP-Seq (GSE51673) | 1e-1968 |
| 10 | [illegible] | Foxf1 / Lung-Foxf1-ChIP-Seq (GSE77951) | 1e-1948 |

**LNCaP/Furamidine – AR ChIP**

| Rank | Motif | Name | p-value |
|---|---|---|---|
| 1 | [illegible] | FOXM1 / MCF7-FOXM1-ChIP-Seq (GSE72977) | 1e-2236 |
| 2 | [illegible] | FOXA1 / LNCAP-FOXA1-ChIP-Seq (GSE27824) | 1e-2191 |
| 3 | [illegible] | FOXA1 / MCF7-FOXA1-ChIP-Seq (GSE26831) | 1e-2186 |
| 4 | [illegible] | Foxa2 / Liver-Foxa2-ChIP-Seq (GSE25694) | 1e-1739 |
| 5 | [illegible] | Fox:Ebox / Panc1-Foxa2-ChIP-Seq (GSE47459) | 1e-1687 |
| 6 | [illegible] | Foxa3 / Liver-Foxa3-ChIP-Seq (GSE77670) | 1e-1486 |
| 7 | [illegible] | FOXA1:AR / LNCAP-AR-ChIP-Seq (GSE27824) | 1e-1408 |
| 8 | [illegible] | GRHL2 / HBE-GRHL2-ChIP-Seq (GSE46194) | 1e-1356 |
| 9 | [illegible] | ARE / LNCAP-AR-ChIP-Seq (GSE27824) | 1e-1047 |
| 10 | [illegible] | FoxL2 / Ovary-FoxL2-ChIP-Seq (GSE60858) | 1e-987 |

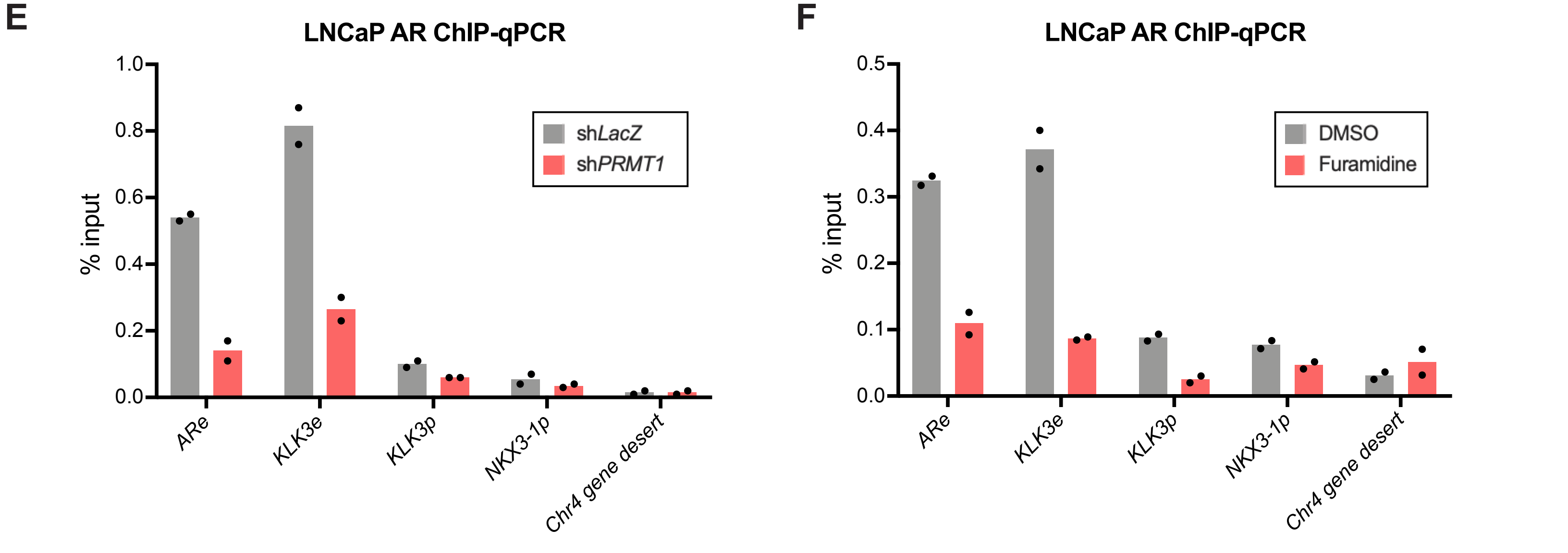

**Figure S4. Additional analyses related to AR ChIP-seq in LNCaP cells. Related to Figure 4.**
(A and B) Pie charts showing the distribution of AR binding sites across the indicated genomic regions in LNCaP cells subjected to control or *PRMT1* knockdown (A) or treated with DMSO or furamidine (B).
(C and D) Motif analysis of genomic regions enriched in AR binding sites in LNCaP cells subjected to control or *PRMT1* knockdown (C) or treated with DMSO or furamidine (D). The top 10 enriched motifs in each condition are shown, ranked by p-value.
(E and F) ChIP-qPCR showing AR enrichment at the *AR* and *KLK3* enhancers (*ARe* and *KLK3e*) and *KLK3* and *NKX3-1* promoters in LNCaP cells expressing sh*LacZ* or sh*PRMT1* (E) or treated with DMSO or furamidine (F). AR enrichment at a control region (*Chr4 gene desert*) is also shown. Data are shown as percentage of ChIP input; bars represent the mean of n = 2 replicates.

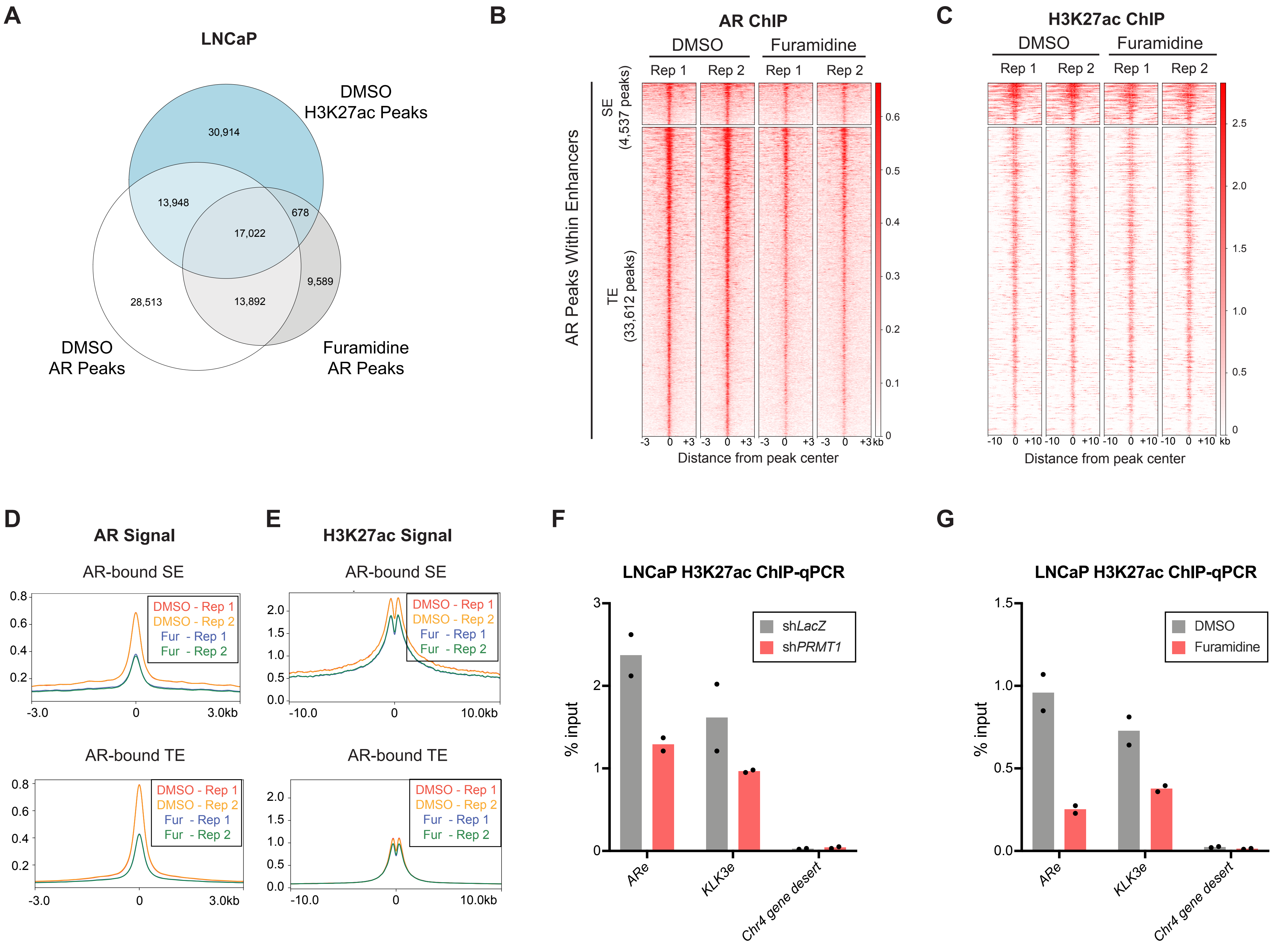

A
LNCaP
DMSO H3K27ac Peaks
30,914
13,948
678
17,022
9,589
28,513
13,892
DMSO AR Peaks
Furamidine AR Peaks
B
AR ChIP
DMSO
Furamidine
Rep 1
Rep 2
Rep 1
Rep 2
AR Peaks Within Enhancers
SE (4,537 peaks)
TE (33,612 peaks)
Distance from peak center
C
H3K27ac ChIP
DMSO
Furamidine
Rep 1
Rep 2
Rep 1
Rep 2
Distance from peak center
D
AR Signal
AR-bound SE
AR-bound TE
DMSO - Rep 1
DMSO - Rep 2
Fur - Rep 1
Fur - Rep 2
E
H3K27ac Signal
AR-bound SE
AR-bound TE
F
LNCaP H3K27ac ChIP-qPCR
% input
shLacZ
shPRMT1
ARe
KLK3e
Chr4 gene desert
G
LNCaP H3K27ac ChIP-qPCR
% input
DMSO
Furamidine
ARe
KLK3e
Chr4 gene desert

**Figure S5. Small-molecule PRMT1 inhibition leads to loss of AR binding and H3K27 acetylation at enhancer regions. Related to Figure 5.**

(A) Venn diagram of overlap between H3K27ac peaks in DMSO-treated LNCaP cells and AR peaks in DMSO- or furamidine-treated LNCaP cells.

(B and C) Heatmaps of AR (B) and H3K27ac (C) ChIP-seq signal over AR peaks in superenhancer (SE) or typical enhancer (TE) regions, shown in the context of DMSO or furamidine treatment. Peaks are rank-ordered by AR signal within 3 kb of the peak center. H3K27ac signal is shown over 10 kb flanking the peak center. Two replicates are shown for each condition.

(D and E) Profile plots of average AR (D) and H3K27ac (E) signal in the regions shown in (B) and (C).

(F and G) ChIP-qPCR validation of decreased H3K27ac enrichment at enhancer elements upstream of *AR* and *KLK3* (*ARe* and *KLK3e*) upon *PRMT1* knockdown (F) or furamidine treatment (G). H3K27ac enrichment at a control region (*Chr4 gene desert*) is also shown. Data are shown as percentage of ChIP input; bars represent the mean of n = 2 replicates.

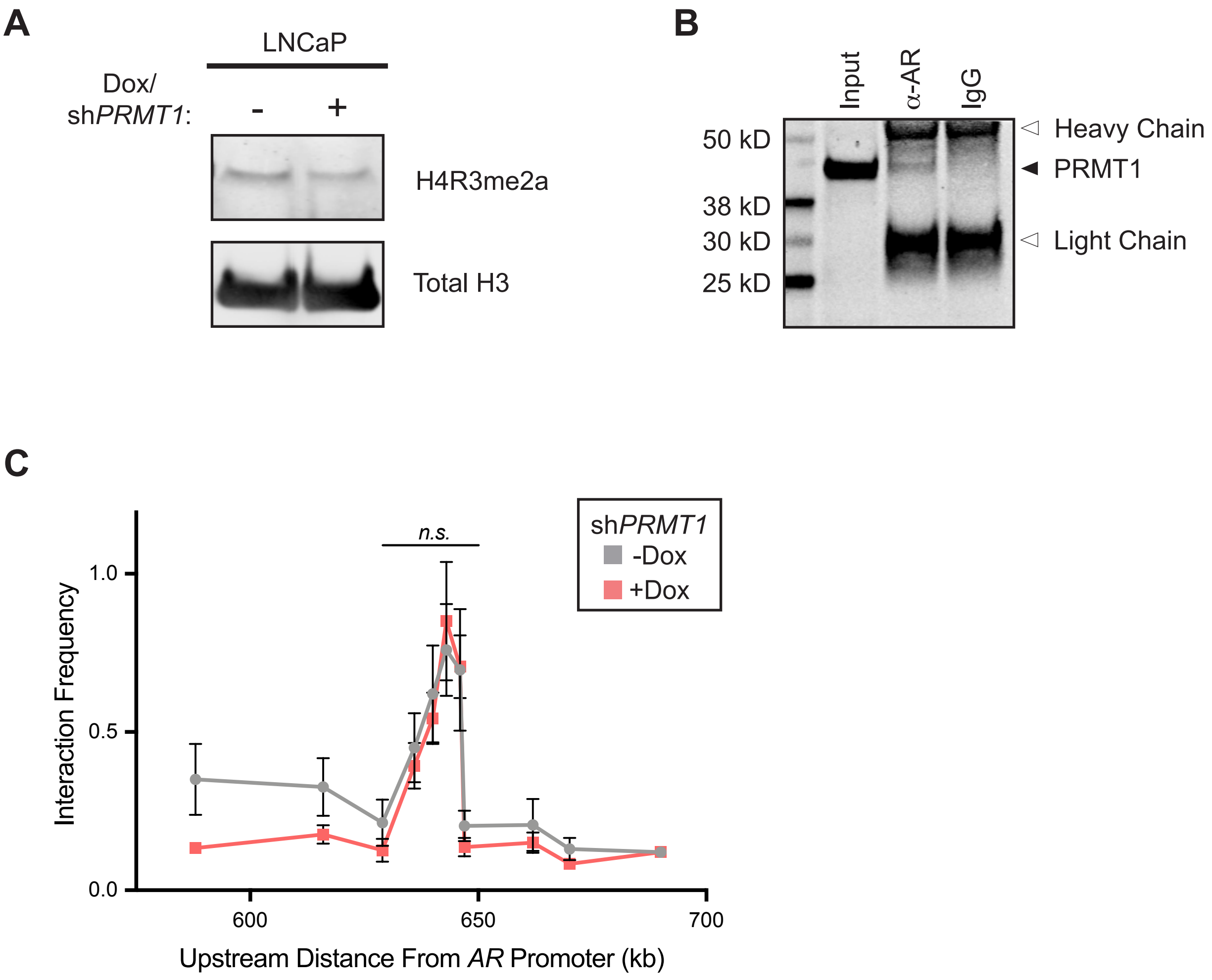


**Figure S6. Role of PRMT1 in modulating AR transcriptional activity. Related to Figure 5.**
(A) Western blot showing decreased H4R3me2a in LNCaP cells upon *PRMT1* knockdown. Histones were isolated for immunoblotting analysis by histone acid extraction.
(B) Co-immunoprecipitation assay showing association of PRMT1 with AR on chromatin in LNCaP cells. Normal rabbit IgG was used as a negative control.
(C) Chromosome conformation capture (3C) experiment showing enhancer-promoter interactions at the *AR* locus in LNCaP cells with or without dox-induced *PRMT1* knockdown. Interaction frequencies were measured by qPCR using the *AR* promoter as bait and are normalized to a BAC spanning the region. Error bars represent mean ± SEM, n = 3 replicates.

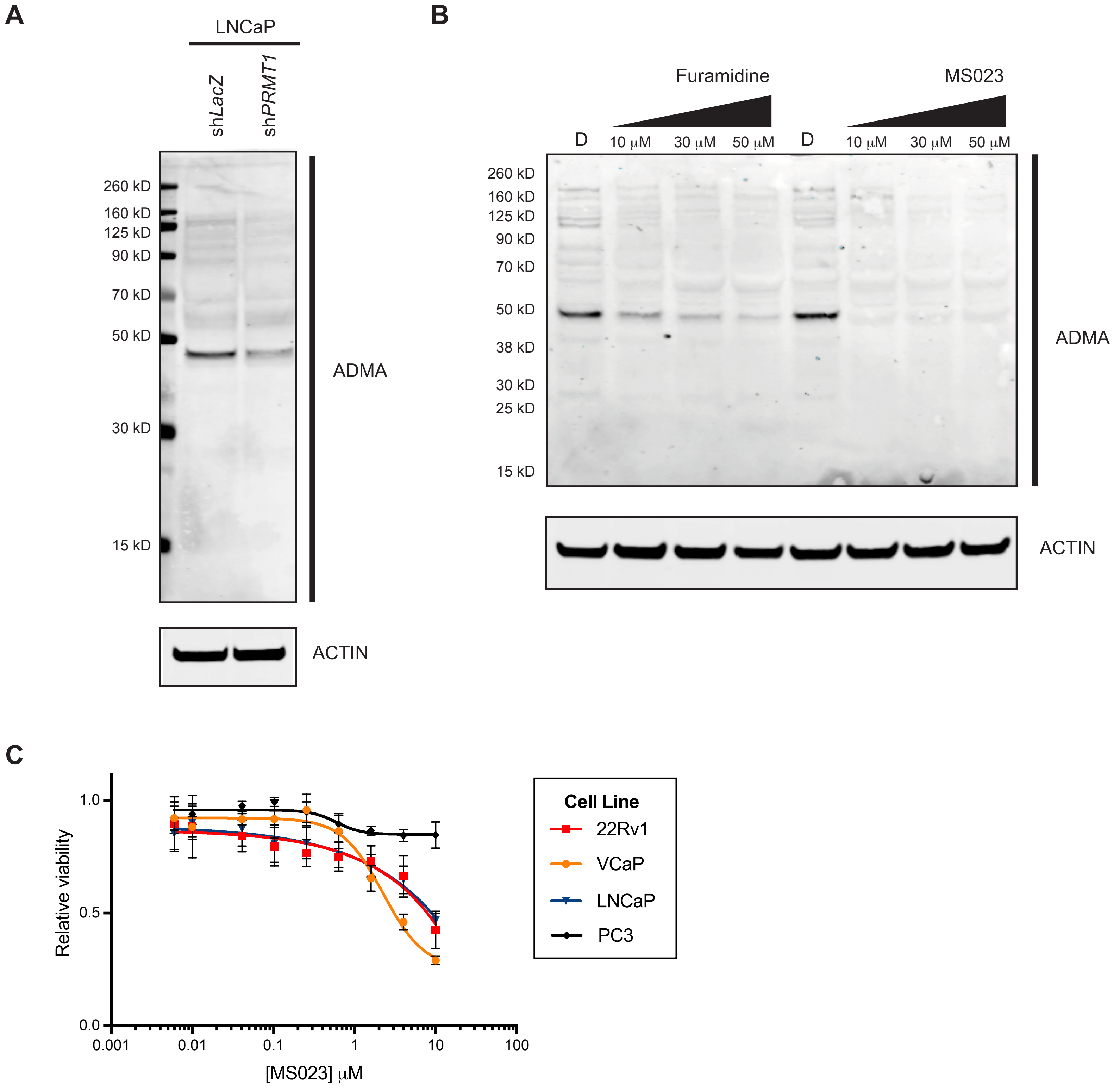


**Figure S7. Validation of *PRMT1* knockdown and PRMT1 inhibitor activity. Related to Figure 6.**
(A) Western blot showing global decrease in asymmetric dimethyl arginine (ADMA) upon *PRMT1* knockdown.
(B) Western blot showing global decreases in ADMA upon treatment with the small-molecule PRMT1 inhibitors furamidine and MS023 compared to the DMSO (D) control.
(C) Relative viability of prostate cancer cell lines after 5 days of treatment with MS023 at the indicated concentrations, normalized to the DMSO condition. Error bars represent mean ± SD, n = 3 replicates.

**Table S7. shRNA and sgRNA target sequences.**

| Name | Target sequence (5' - 3') | Name | Target sequence (5' - 3') |
|---|---|---|---|
| shLacZ | TCGTATTACAACGTCGTGACT | POLR3H_sg1 | GCTCAACGACTCCATTGCCG |
| shPRMT1 | GCCTACTTCAACATCGAGTTC | POLR3H_sg2 | ATGCAGGTCGTGTACAACGT |
| shAREx1 | GAGCACTGAAGATACTGCTGAGTAT | POLR3H_sg3 | GCCATCCCCAGGGAATACAT |
| shAR-V7 | GTAGTTGTGAGTATCATGA | POLR3K_sg1 | GGCACGTGTTGCAGGCGAAG |
| shAREx7 | TCAAGGAACTCGATCGTAT | POLR3K_sg2 | GCACAACATCACCCGCAAGG |
| API5_sg1 | CTGGGGTAGTCAAGGTACCG | POLR3K_sg3 | GGGTGATGTTGTGCACGTAG |
| API5_sg2 | TGCACAGTTAGACCTCTGTG | PRMT1_sg1 | GGGTCCACGACATCCACTAG |
| API5_sg3 | GCCTATCAAGTGATATTGGA | PRMT1_sg2 | GATGGCCGTCACATACAGCG |
| AR_sg1 | AGGGTACCACACATCAGGTG | PRMT1_sg3 | AAAGCCAACAAGTTAGACCA |
| AR_sg2 | GGACGCAACCTCTCTCGGGG | PRMT5_sg1 | GGAGAAAAACCCAAATGCCG |
| AR_sg3 | CCTTAAAGACATCCTGAGCG | PRMT5_sg2 | TGCACCAACTACACACACAG |
| CHTOP_sg1 | AGGGGCGGGATGTCACTCCG | PRMT5_sg3 | GAAGATTCGCAGGAACTCCG |
| CHTOP_sg2 | CCAAGATGTCTCTAAATGAG | PRPF4B_sg1 | GAAAACGACGAGAACCAGAG |
| CHTOP_sg3 | ACGGTTAGGCCGACCCATAG | PRPF4B_sg2 | TTTACCTCTTAAATCAACTG |
| DBR1_sg1 | GCTGTTACACTAAAGTTACA | PRPF4B_sg3 | ATTATGCTTGGCTTTCACTG |
| DBR1_sg2 | AGGCGGCAAACTTCACATGA | SART1_sg1 | GCCGTCGGACGACACCCGAG |
| DBR1_sg3 | GCTGGTGTGGTAAAATACCG | SART1_sg2 | GAAGCGCGATGACGGCTACG |
| EFTUD2_sg1 | GCTTGGCATCCACCTGACGA | SART1_sg3 | CTCCGAATACCTCACGCCTG |
| EFTUD2_sg2 | AGCCACATGCCCTTACATCA | SFPQ_sg1 | ATGATCGTGGAAGATCTACA |
| EFTUD2_sg3 | GAACACGGTTCACCTCGATG | SFPQ_sg2 | ATGGGCCTCAATCAGAATCG |
| NUDT21_sg1 | AGCCAGATTTCAGCGCATGA | SFPQ_sg3 | TCTACCTGCTGATATCACGG |
| NUDT21_sg2 | CCTGGTGGTGAACTTAACCC | sgControl98 | ATCGTTTCCGCTTAACGGCG |
| NUDT21_sg3 | ACTAAAACGCTTAATGACAG | sgGFP | GAAGTTCGAGGGCGACACCC |

**Table S8. Primer sequences for qPCR and genotyping assays.**

| Name | Forward primer (5' - 3') | Reverse primer (5' - 3') |
|---|---|---|
| AR-FL RT-qPCR | CAGCCTATTGCGAGAGAGCTG | GAAAGGATCTTGGGCACTTGC |
| AR-V7 RT-qPCR | CCATCTTGTCGTCTTCGGAAATGTTA | TTTGAATGAGGCAAGTCAGCCTTTCT |
| KLK3 RT-qPCR | GAGCAGCCCTATCAACCCCCTATT | AGCAACCCTGGACCTCACACCTAA |
| PRMT1 RT-qPCR | GGAAAGCAGTGAGAAGCCCA | CGTCCTTCAGCATCTCCTCG |
| GAPDH RT-qPCR | GTCTCCTCTGACTTCAACAGCG | ACCACCCTGTTGCTGTAGCCAA |
| ACTB RT-qPCR | AGAGCTACGAGCTGCCTGAC | AGCACTGTGTTGGCGTACAG |
| ARe ChIP-qPCR | CCAGACAGGCAAGCTTTCAG | TGGGCAACTTCCAATGACTG |
| KLK3e ChIP-qPCR | TGGGACAACTTGCAAACCTG | CCAGAGTAGGTCTGTTTTCAATCCA |
| KLK3p ChIP-qPCR | CCTAGATGAAGTCTCCATGAGCTACA | GGGAGGGAGAGCTAGCACTTG |
| NKX3-1p ChIP-qPCR | GCAGATCTGAGTTTGCACCA | TGGGACGATCAAGACAAACA |
| Chr4 gene desert ChIP-qPCR | Human Negative Control Primer Set 2 (Active Motif, #71002) | |
| AR-V7-GFP genotyping | CCAACTTTACATGCTGCTTCC | TGCTTGTCGGCCATGATATAG |
| AR-V7 genotyping | TCAGGTTCCATTCTTCTCAGTCCAGTT | GGTCTGGTCATTTTGAGATGC |
